# HIV-1 infection of human microglia activates inflammatory pathways associated with HIV-associated neurocognitive disorders (HAND)

**DOI:** 10.1101/2025.02.13.638028

**Authors:** Jason E. Hammonds, Roy Moscona, Kathleen Candor, Hope Guthier, Rathnakumar Kumaragurubaran, Thomas Hagan, Paul Spearman

**Affiliations:** Division of Infectious Diseases, Department of Pediatrics, Cincinnati Children’s Hospital and University of Cincinnati, Cincinnati, OH, USA; Division of Rheumatology, Department of Medicine, University of Pennsylvania Perelman School of Medicine, Philadelphia, PA, USA; Pathobiology & Molecular Medicine Graduate Program, University of Cincinnati and Cincinnati Children’s Hospital, Cincinnati, OH, USA; Division of Developmental Biology, Single Cell Genomics Core, Department of Pediatrics, Cincinnati Children’s Hospital and University of Cincinnati, Cincinnati, OH, USA

**Keywords:** HIV-1, microglia, HIV-associated neurocognitive disorders, HAND, neuropathogenesis

## Abstract

HIV-associated neurocognitive disorders (HAND) cause significant dysfunction among people living with HIV. Microglia are the primary immune cells of the central nervous system and are readily infected by HIV. Microglia are thought to contribute to neuroinflammation and cognitive dysfunction in neurodegenerative diseases such as Alzheimer’s Disease and are likely to play an important role in the pathogenesis of HAND. In order to identify pathways that may contribute to neuropathogenesis in HAND, we infected induced pluripotent stem cell-derived microglia (iMG) with HIV and defined gene expression changes over an 8-day period. Monocyte-derived macrophages (MDMs) were studied in parallel to identify common pathways stimulated in myeloid cells versus the unique aspects of microglia infection. Infection of iMG led to the induction of a robust early inflammatory response triggered within hours of infection, a pattern that differed significantly from that seen in MDMs. Remarkably, gene expression changes in iMG reproduced many of the characteristic genetic signatures previously identified in brain tissues obtained from individuals clinically diagnosed with HAND. Inflammatory activation representing interferon-mediated signaling, TNF/NF-κB, and IL-6/JAK/STAT pathways were particularly prominent over the time course of infection. Interferon-mediated signaling led to enhanced expression of multiple HIV restriction factors, yet viral replication in iMG remained robust. These findings suggest that HIV infection of microglia is the key cellular driver of neuroinflammation in the CNS of HIV-infected individuals. Further studies of infected microglia are likely to aid in understanding the pathogenesis of HAND and in evaluating therapeutic strategies to limit or eliminate HIV-induced neuropathogenesis.

**Author Summary:** Persons living with HIV frequently develop debilitating problems with brain function that are collectively known as HIV-associated neurocognitive disorders (HAND). Treatment with antiretroviral drugs to control HIV has largely eliminated the most severe form of HIV-related brain dysfunction, HIV encephalitis, but a significant proportion of HIV-infected individuals receiving antiviral therapy still develop significant problems with cognitive function. Microglia are the major macrophage-like cells of the brain and are highly susceptible to HIV infection. Inflammation in the brain following infection contributes to damage to neurons and is the likely source of decline of brain function. This study used microglia that were derived from induced pluripotent stem cells, termed iMG, to study the kinetics of gene expression changes in HIV-infected microglia and compared those changes to those seen in monocyte-derived macrophages (MDMs). iMG were found to be easily infected with a macrophage-tropic HIV strain and exhibited a rapid increase in gene expression associated with inflammatory signaling through interferon- and tumor necrosis factor/NF-kappa B-mediated pathways. Remarkably, the patterns of gene expression in microglia strongly overlapped with gene expression signatures derived from brain tissue samples of HIV patients with HAND. The rapid onset and magnitude of inflammation as well as induction of specific HIV restriction factors differed from that seen in infected macrophages/MDMs. These studies reinforce the central role of microglia in the pathogenesis of HAND and provide insights into HIV-microglia interactions that can help direct interventions to reduce inflammation and preserve brain function.

## Introduction

HIV-1 infection remains a significant global health challenge, with an estimated 40.8 million people living with HIV-1 (PLWH) as of 2024 [1]. The availability and utilization of combination antiretroviral therapy (cART) have significantly lowered the morbidity and mortality associated with HIV infection, shifting HIV-1 infection from a fatal illness to a manageable chronic condition [2–5]. Despite this remarkable advance, cART fails to eradicate long-term tissue reservoirs harboring infectious virus, and HIV-related comorbidities remain a problem even in individuals who successfully control their plasma viral load. The central nervous system (CNS) has distinct biological and pharmacological properties that differentiate it from other tissues [6]. An important characteristic is the brain parenchyma’s relative isolation from peripheral immune surveillance. Immune defense within the parenchyma is primarily managed by microglia, specialized myeloid cells that reside permanently within brain tissue [7]. Myeloid cells, including microglia, are susceptible to and permissive for HIV replication, and contribute to tissue reservoirs for the virus [8–14]. HIV infects microglia in the brain at early times following acute infection, likely through infected peripheral blood monocytes and lymphocytes that cross the blood-brain barrier [15]. Following infection, microglia serve as a source of ongoing HIV replication and spread within the CNS, contributing to inflammation and neurodegeneration, in addition to serving as a long-lived latent reservoir.

HIV infection of microglia and resident brain macrophages contributes to neurologic and neurocognitive complications, spanning from subtle neurocognitive difficulties to severe, debilitating dementia [16–18]. Collectively, these conditions are classified as HIV-associated neurocognitive disorders (HAND). The development of cART has significantly lowered the prevalence of HIV-associated dementia (HAD); however, its impact on the other primary forms of HIV-associated neurocognitive disorders (HAND), asymptomatic neurocognitive impairment (ANI) and mild neurocognitive disorder (MND), has been more limited [3, 19–21]. Even with access to cART, an estimated 20-50% of HIV-infected individuals experience some degree of neurocognitive impairment [22–24]. Microglia normally perform neuroprotective roles by clearing apoptotic cells and releasing neurotrophic factors and growth hormones that support cellular health in brain parenchyma. In contrast, persistent inflammation imparted by chronically activated microglia has been shown to affect neural plasticity and memory degradation, contributing to the pathogenesis of many neurodegenerative disorders [25, 26]. In cases of neuroinflammation and neurodegeneration, chronically activated microglia release a variety of pro-inflammatory cytokines, including tumor necrosis factor-α (TNF-α), interleukin-6 (IL-6), interleukin-1β (IL-1β), reactive oxygen species (ROS), and chemokines such as CXCL10 and MCP-1 [24]. The HAND signature derived from brains of PLWH shares many features with other chronic neurocognitive diseases, including Alzheimer’s disease and multiple sclerosis [19, 21].

Research on HIV-related neuropathology in humans has largely been restricted to analyzing brain tissues obtained postmortem. Due to sample degradation prior to collection, high-resolution analyses of microglia activation in response to HIV-infection have not been possible from infected brain tissues. To study in detail the response of microglia to HIV infection, we generated and infected induced pluripotent stem-derived microglia (iMG) [27–29] with an M-tropic HIV-1 primary isolate. To provide context, parallel analyses were performed using human whole-blood isolated, primary monocyte-derived macrophages (MDMs). We found marked differences between inflammatory pathway activation induced in iMG vs. MDMs, with an initial response in iMG that was marked by significant type 1 IFN, TNF, IL-6 and IL-1β production, and characterized by a broader and more potent induction of inflammatory pathways over time than that seen in MDMs. The signature elicited in iMG matched remarkably well with the signatures of HAND previously characterized from brains of subjects exhibiting this clinical disorder.

## Results

### Gene expression analysis of human iPSC-derived microglia using signature profiles from primary human microglia

Human microglia (iMG) were generated following well-established methods first described by the Blurton-Jones laboratory and adapted by our laboratory [27–29]. Initial validation of iMG in our laboratory as a relevant model for HIV studies involved assessment of phenotypic marker expression using flow cytometry and immunofluorescence microscopy. iMG expressed characteristic myeloid and microglial markers, including CD45, CD11b and CX3CR1, as well as HIV-entry receptors CD4, CCR5 and CXCR4 (S1A Fig). Immunofluorescence further confirmed expression of microglial lineage markers P2RY12, IBA1, PU.1 and CX3CR1, with characteristic morphology visible by DIC imaging (S1B Fig). These phenotypic features reflect those of human microglia [27, 30–32] and support the use of iMG as a biologically relevant model for HIV–microglia infection studies. Gene expression in iMG was quantified via RNA sequencing and compared against a core transcriptomic profile of primary human adult microglia (AMG) to confirm its relevance as a robust human microglia model. As an initial step, we performed principal component analysis (PCA) of gene expression from iMG alongside AMG [30], homeostatic adult microglia from the Human Microglia Atlas (HuMicA_HOM) [33], CD14+ monocytes (CD14M) [29], MDMs, and an alternative iMG population from the Blurton-Jones laboratory (labelled as iMG_Abud) [27, 28]. In this analysis, both sources of iMG exhibited greater similarity to AMG than the two primary myeloid cell types, MDMs and CD14+ monocytes, across both principal axes of variance (Fig 1A). The previously established iMG-Abud gene expression profile clustered with the iMG from the current study, indicating similarity in the transcriptional profiles of the distinct iMG lines. To further analyze similarities and differences in gene expression between microglia and macrophages, gene-level fold changes between both iMG and AMG against MDM were computed (Fig 1B). Among genes highly differentially expressed in adult microglia compared to macrophages, there was a strong correlation between expression in *ex vivo* brain-derived microglia and iMGs (R = 0.47, p < 2.2e−16), indicating a common transcriptional profile that is distinct from MDMs. The top right quadrant in Fig 1B represents genes upregulated in both types of microglia, while the bottom left quadrant shows genes upregulated in MDMs. Genes highly expressed in both iMG and AMG relative to MDMs include those identified in several signature human microglia studies, including CX3CR1, OLFML3, SYT6, ACY3, SMARCA1, TAL1, WNT5A, and FCGBP (Fig 1B) [30, 31, 34–37]. Despite the shared myeloid lineage of microglia and MDMs, comparative analysis here identified 30 distinct marker genes uniquely or highly differentially expressed (top 30, log_2_FC≥7) in microglia (Fig 1C). Hallmark microglia signature genes including P2RY12, OLFML3, CX3CR1, and ADGRG1 were differentially expressed as expected, yet these genes also had a detectable level of expression in MDMs (Fig 1C, genes shown in cluster 1). Additionally, we examined baseline expression levels of a panel of core microglial signature genes across different microglial populations (iMG, AMG, HuMicA) and monocyte-derived macrophages (MDMs) (S2 Fig). While many of these genes were also expressed in MDMs albeit often at significantly lower levels, the overall gene expression profile was largely conserved between iMG and adult human microglia. In contrast, we identified novel microglia markers that were distinct from MDMs, including genes related to GO ontology categories of somatodendritic compartment (CACNA1D, KCNQ3, SYNDIG1, PCLO, IGF1, GABRA3, CTTNBP2, ENPP1, CX3CR1), microglial cell migration (P2RY12, ENPP1, CX3CR1) and gliogenesis (LYVE1, P2RY12, WNT5A, CXCL12, CX3CR1) (Fig 1C, cluster 2). These microglia-specific genes are not present in the well-established microglia signatures reported to date and may represent useful markers for future studies [30, 31, 35]. To better characterize differential gene expression common to microglia versus MDMs, microglia genes with high expression (Fig 1B scatterplot quadrant 1 genes with log_2_FC>2 as compared to uninfected cells), were subjected to over representation analysis (ORA). This analysis revealed several shared enriched GO ontological biological processes (BPs) associated with CNS development and immune function for microglia, which fell into four functional categories: synaptic activity, nervous system development, immune response, and cellular adhesion (Fig 1D). These results align with findings from prior foundational studies performed from *ex vivo* human microglia [30, 31, 38, 39]. Taken together, these analyses support the authenticity of iMG as a model system that can be employed for studying HIV interactions.

**Fig 1.**
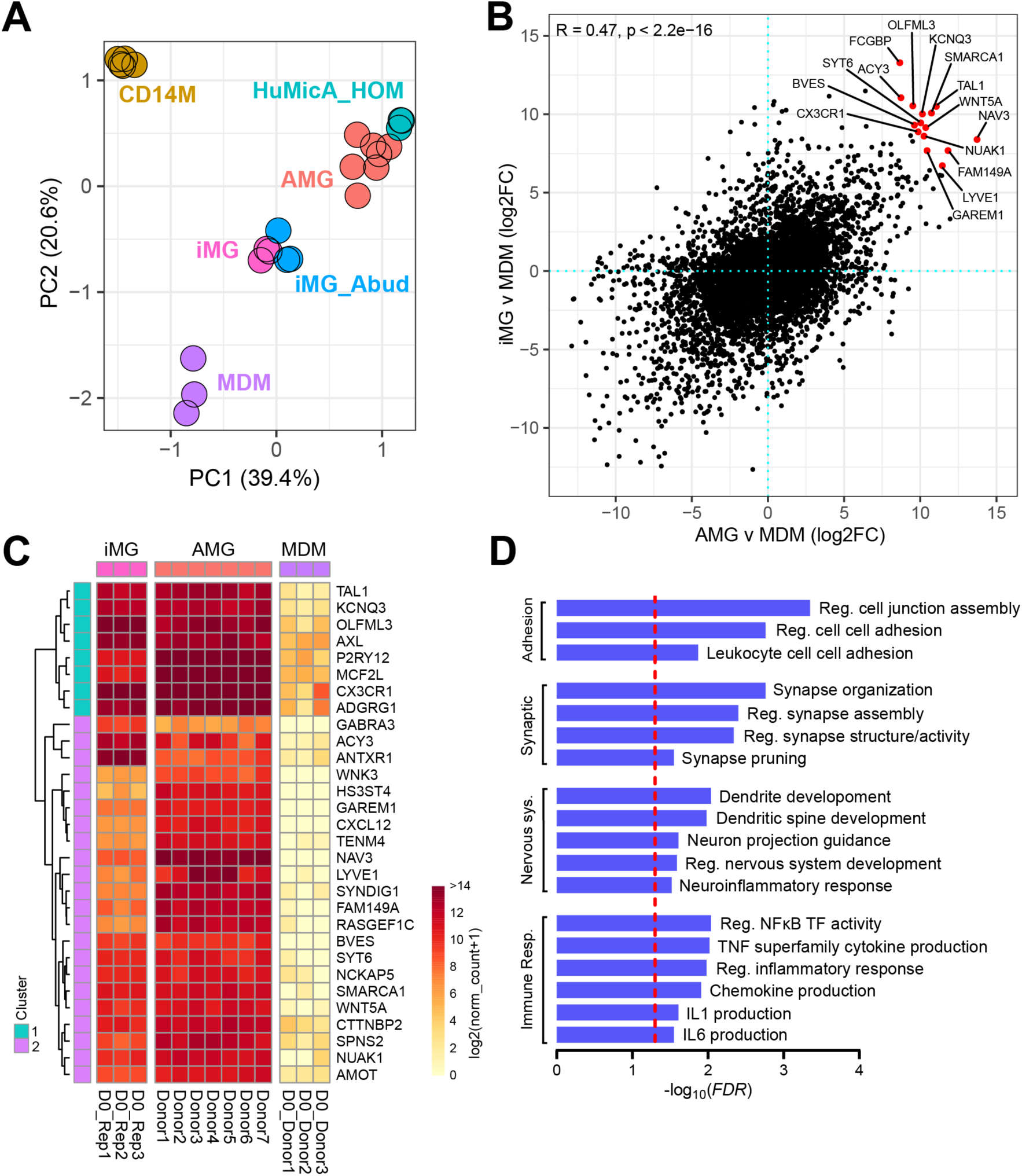
iMG resemble primary human microglia and exhibit a transcriptomic signature distinct from human MDMs. **(A)** Principal component analysis utilizing a core gene expression signature of human microglia [35]. iMG (pink, n=3 biological replicates), iMG_Abud (green) [27], human adult microglia (AMG, peach) [30], homeostatic adult microglia from the Human Microglia Atlas (HuMicA, green) [33], human monocyte-derived macrophage (MDM, purple, n=3 donors from this study), and CD14 monocytes (CD14M, olive green) [28]. **(B)** Scatter plot of gene expression fold changes between iMG vs MDM and AMG vs MDM. Correlation coefficient shown for genes expressed in either AMG or iMG vs MDM. **(C)** Heatmap of top 30 microglia marker genes identified by differential expression analysis between human adult microglia [30], iMG, and MDMs from our study. Microglia signature genes were defined as commonly differentially expressed (FDR<0.05) between both microglia cell types (AMG and iMG) and MDMs (FDR<0.05, absolute log_2_FC≥7, MDM norm_counts<50). **(D)** GO-BP ontology showing enrichment of genes belonging to Q1 of panel B scatterplot, meeting criteria of log2FC>2 for both comparisons (iMG vs MDM and AMG vs MDM). Bars represent a significant level of FDR values. Red dashed line represents the significant threshold (FDR<0.05).

### Response of iMGs to HIV-1 Infection

The results above establish a high transcriptional similarity between the iMG model and primary human microglia and distinguish both from the basal transcriptional profile of MDMs. We next sought to comprehensively define differences in gene expression between iMG and MDMs following HIV-1 infection. To establish the infection model, iMG and MDMs were infected with the R5 M-tropic HIV-1 isolate BaL at a multiplicity of infection (MOI) ranging from 0.05 to 0.5, and viral production monitored by measurement of supernatant p24 antigen over time. Infection was readily established in iMG, even at the lowest MOI (Fig 2A). A consistent release of virus was observed, representing continuous, low-level release commonly exhibited by HIV-1-infected myeloid cell populations. This trend closely mirrored the pattern observed in human MDMs infected with HIV-1 BaL (Fig 2B). Following growth curve analysis, flow cytometry was performed to provide cellular-resolution quantification of infection kinetics in iMG (S3 Fig). Quantification of intracellular p24 demonstrated a low level of infection early after exposure to HIV BaL, with mean infection levels of 2.8% at day 1 and 3.4% at day 2. Infection increased substantially by day 5 (21.4%) and rose sharply thereafter, reaching 77.8% p24⁺ cells by day 8 (n = 3 biological replicates). These data provide refined single-cell validation of infection dynamics and demonstrate efficient viral propagation in iMG. For comparison of changes in gene expression for the remainder of these studies, an MOI of 0.25 was chosen as the input, with RNA samples harvested on days 0, 1, 2, 4, 6, and 8 for comprehensive gene expression analysis. The study of gene expression was concluded on day 8 in order to minimize effects resulting from virus-induced cytotoxicity that occurred at later timepoints [28]. As an additional readout of replication, sample reads that mapped to the HIV-1 BaL genome (GenBank: AY713409.1) were quantified. Logarithmic increases in normalized *Gag* and *Env* counts were observed between days 2 and 8 (Fig 2C and 2D), indicating comparable transcription and ongoing replication in iMG and MDMs.

**Fig 2.**
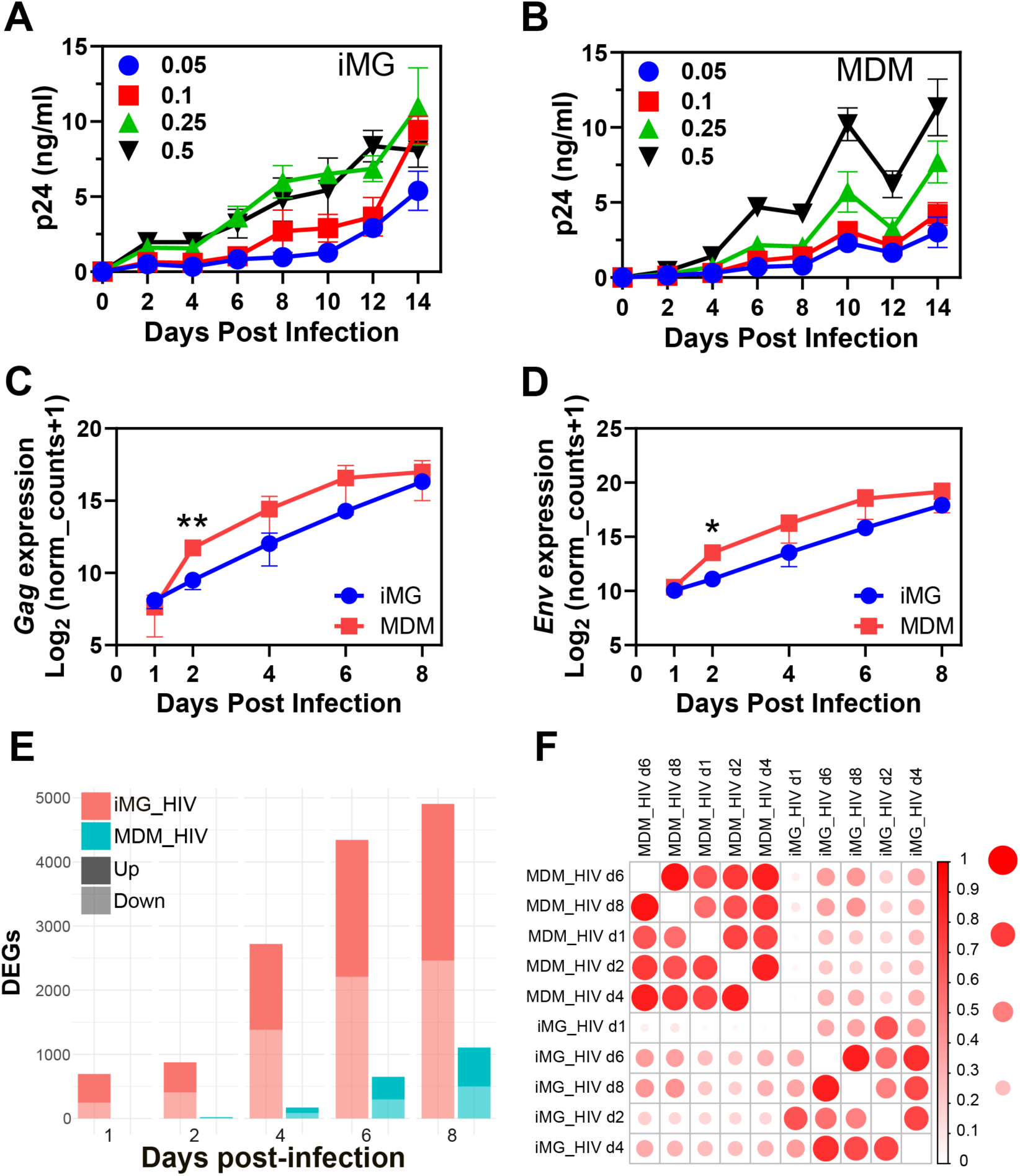
HIV replication drives increasing transcriptional changes in iMGs and MDMs, while iMGs display a distinct gene expression profile. (**A)** HIV BaL growth curve spanning a log range of MOIs in iMG as measured by p24 ELISA from harvested cell culture supernatants. iMG cultures were fed half-media changes every 2 days throughout growth curve. **(B)** HIV BaL growth curve at a range of MOIs from MDMs. MDM culture media was replenished completely on days 3, 6 and 10 post-infection. (**C)** Number of reads mapped to *gag* from the HIV BaL genome in iMG and MDMs over the experimental time course. Statistical significance determined using two-tailed, standard, unpaired t-test. Single timepoint comparing values between iMG and MDMs on day 2 post-infection was significant (p≤0.01, **) while all other time points post-infection showed no significant difference between iMG and MDMs (p>0.05). **(D)** Number of reads mapped to *env* in iMG and MDMs. Single timepoint comparing values between iMG and MDMs on day 2 post-infection was significant (p≤0.05, *) while all other time points post-infection showed no significant differences (p>0.05). **(E)** Number of differentially expressed genes in iMG and MDMs during the 8-day infection time course. **(F)** Spearman correlation matrix comparing gene-level fold changes in iMG and MDMs during HIV infection time course. Statistical significance determined using two-tailed, standard, unpaired t-test.

Global gene expression analysis of HIV-infected iMGs compared to uninfected controls demonstrated an increasing number of differentially expressed genes (DEGs) throughout the infection time course (Fig 2E). The total number of infection-induced DEGs in iMG rose from 629 on day 1 to 4719 on day 8. In contrast, the number of DEGs for HIV-infected MDMs was markedly lower, with no DEGs identified on day 1 and a peak of 1060 on day 8 post-infection. Additionally, principal component analysis of global gene expression in HIV-infected iMGs and MDMs over the 8-day time course revealed that cell-type differences were the primary source of variation, as indicated by separation along PC1. However, infection-driven expression changes were conserved between the two cell types, as shown by the parallel trajectories along PC2 throughout the time course (S4A Fig). As MDMs from repeated experiments represent primary cells derived from multiple donors, while iMG are isogenic, the lower number of DEGs identified in infected MDMs could simply result from increased variance in responses due to genetic diversity. To investigate this, we examined the variance and fold changes of infection-induced DEGs identified from iMG in both cell types. We found that while MDMs did have increased variance in their response (S4B Fig), they also exhibited decreased fold changes in these genes compared to iMG (S4C Fig). This indicates that the difference in DEGs was not solely due to genetic variance in MDMs, but that instead HIV triggers a more robust transcriptional response in iMG compared to MDMs. We next compared the similarity in responses across cell types and time points post HIV-infection by generating a correlation matrix, allowing pairwise comparisons of gene-level infection-induced fold changes. The infection correlation matrix generated distinct clusters specific for iMG and MDMs (Fig 2F). The fold changes in iMG on day 1 (iMG_HIV d1) did not correlate with those from MDMs at any time point, indicating that responses on day one post-infection were unique to iMGs. The highest correlations between cell types occurred on days 6 and 8 post-infection, suggesting a more similar response at later time points. Furthermore, responses were largely cell type-dependent, rather than demonstrating a conserved temporal response to HIV infection common to both iMG and MDMs. Together, these findings establish that following infection with HIV, viral replication is similar in iMG and MDM cultures, while infection of iMG generates an accelerated transcriptional response that is of higher magnitude and distinct from that seen in MDMs.

### Genome wide RNA expression analysis of HIV-infected iMG and MDMs

The remarkable differences in gene expression between iMG and MDMs following HIV infection were next examined via gene set enrichment analysis (GSEA) using the Hallmark gene set collection from the Molecular Signatures Database (MSigDB) [40, 41] (Fig 3A). This analysis highlights the early upregulation of inflammatory pathways that is present on day 1 post-infection in iMG and absent in MDMs. Infected iMG upregulated pathways related to IFN responses, TNFα signaling via NF-κB, IL-6/Jak/Stat3 signaling, and inflammatory and complement responses. The early inflammatory response in iMG seen for TNFα, IL-6/JAK/STAT3, and inflammatory response modules then diminished by day 2 before returning on day 6 or 8 (Fig 3A), suggesting a two-phase response to infection. In contrast, MDMs infected with HIV exhibited no positively enriched gene sets on day 1 post-infection, and their later responses (days 6 and 8) were primarily restricted to interferon signaling pathways. Among the downregulated pathways, it was notable that genes related to cell cycle, including E2F targets and G2M checkpoint, were significantly downregulated at all time points following infection in both iMG and in MDMs. To ensure that the early responses in iMG seen here were not an artifact of the iPSC line chosen for iMG development, a parallel comprehensive analysis using iMG derived from a second iPSC clone, referred to as iMG2, was performed. HIV-infection of iMG2 generated responses that were consistent with those observed in iMG by Hallmark gene set enrichment analysis (S5A Fig) and exhibited broad initial and later temporal upregulation of inflammatory genes consistent with similar kinetic changes that had been observed in iMG (S5B Fig). We conclude that the early inflammatory response in iMG, which differs markedly from inflammatory pathway changes in MDMs, is consistent across a second iPSC source and not merely due to cell selection. Furthermore, aging of cultures did not account for the observed changes. We compared DEGs in infected vs. uninfected iMG or MDM cultures of the same age, and there were no significant differences noted (S6 Fig).

**Fig 3.**
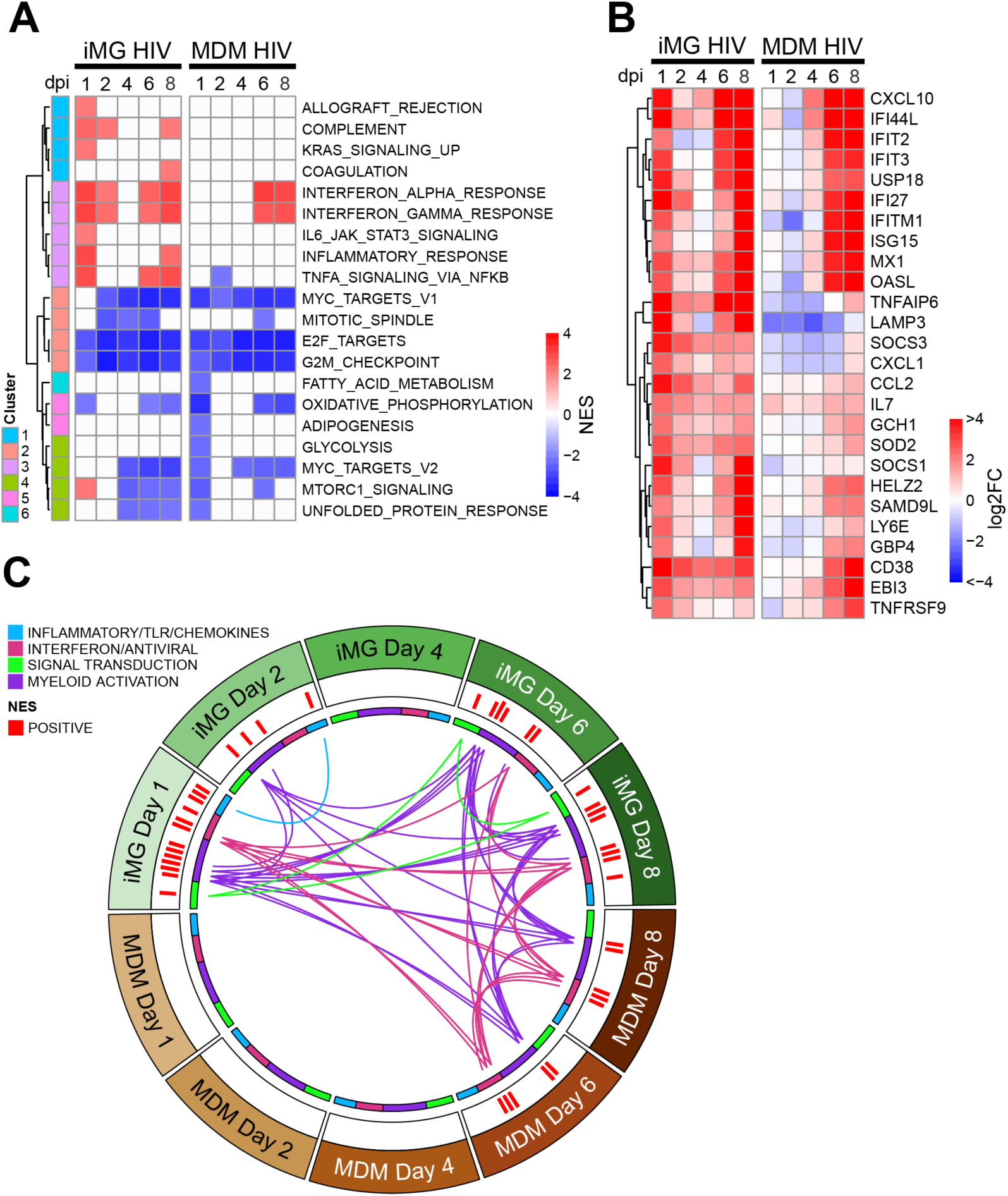
Temporal gene set enrichment analysis demonstrates an early induction of inflammatory responses in HIV-infected iMG. **(A)** Hallmark gene set enrichment analysis of HIV-infected iMG and MDM. Color represents normalized enrichment score (NES) in significantly enriched gene sets (FDR<0.05, absolute NES>2). **(B)** Temporal changes in expression of the most common genes (included in ≥ 25% of gene sets, log_2_ FC≥1) from gene sets in cluster 3 of the Hallmark pathway enrichment analysis (panel A). **(C)** Circos plot of overlap in enriched blood transcriptional modules in HIV-infected iMG and MDMs. Each segment of the circle represents responses in either iMG or MDMs at a single timepoint post-infection. Bars in the outer ring represent significantly enriched modules (FDR<0.05, absolute NES≥2). Lines connect shared modules exhibiting positive enrichment (NES≥2) in more than one cell type/time point. Inner circle and line colors represent the module functional groupings. The blood transcriptional modules were derived from large-scale network integration of publicly available human blood transcriptomes [42, 43].

To conduct a more detailed analysis of the early and late inflammatory responses at the gene level, we examined expression changes over the infection time course of genes enriched in two or more pathways from cluster 3 in Fig 3A, and exhibiting an induction of log_2_ FC>1 in iMG (Fig 3B). Infected iMG broadly upregulated IFN-stimulated genes (ISGs) including IFITM1, USP18, ISG15, IFI27, IFI44L, IFIT2, IFIT3, MX1, OASL, SOCS1 AND SOCS3 on day 1 (Fig 3B). HIV infection of iMG also induced heightened expression of inflammatory chemokine genes (CCL2, CXCL1, CXCL10) together with multiple genes regulated by TNFα/NF-κB signaling (LAMP3, SOD2, TNFRSF9 AND TNFAIP6). On day 1, iMG genes with the most significant log_2_ FC increase in gene expression were TNFAIP6 (log_2_ FC 5.6), IFI44L (4.7), CD38 (4.7), IFI27 (4.6), and LAMP3 (4.3); while 26 additional genes achieved a log_2_ FC of 2 or greater. On day 8, 21 iMG genes achieved gene expression changes exceeding log_2_ FC of ≥2, with the greatest changes observed in CXCL10 (log_2_ FC 9.5), IFI44L (8.6), IFIT2 (7.6), USP18 (7.1) and OASL (6.5). In contrast, HIV infection of MDMs resulted in only a single gene with a log_2_ FC of 1 on day 1 post-infection (IL7, log_2_ FC = 1.0) and 16 genes that achieved a log_2_ FC of ≥2 on day 8 from the same cluster of hallmark pathways. By day 6-8, however, MDMs demonstrated upregulation of a number of IFN-related genes and TNFα/NF-κB-related genes (Fig 3B, MDM heatmap). Together, these results demonstrate that HIV induces significant inflammatory responses in iMG that occur earlier in the time course and are greater in terms of both magnitude and breadth when compared to MDMs, highlighted by ISGs, inflammatory chemokines, and TNFα/NF-κB signaling. To further depict HIV-dependent temporal transcriptome changes in iMG and MDM, we next performed pathway enrichment using an alternative set of widely-used gene modules from human blood transcriptome datasets, known as blood transcriptional modules (BTMs) [42, 43], and examined the overlap across cell types and timepoints via Circos plot (Fig 3C). The connections drawn represent only the upregulated modules/pathways, showing shared modules between iMG and MDMs following infection. Upregulated modules were found in common for iMG at day 1 and 2, with increased commonality between day 1 and days 6 and 8. A unique response in iMG is shown in green (signal transduction) and is in common between days 6 and 8. In contrast, no modules were shared between iMG and MDMs until day 6. Common pathways from this set of modules between infected iMG and MDMs on day 6 and day 8 included IFN/antiviral and myeloid activation pathways (pink and purple modules/connecting lines, Fig 3C). Consistent with the hallmark GSEA, these results illustrate the early and robust hyperinflammatory reactivity of HIV-infected iMG, with a shared late response with MDMs highlighted by IFN-related pathways and pathways indicating myeloid activation.

To determine whether the early inflammatory response observed at day 1 post-HIV infection was confined to productively infected microglia or was also present in uninfected bystander cells, we performed single cell RNA-sequencing (scRNA-seq) and analyzed cells stratified by viral RNA abundance (S7 Fig). Temporal transcriptional shifts (S7A Fig) tracked with increasing HIV *pol* RNA abundance (S7B Fig), indicating HIV-driven transcriptional remodeling. Consistent with the early pathway activation observed in bulk RNA-seq (Fig 3), gene set enrichment analysis of scRNA-seq pseudobulk profiles showed highly concordant enrichment of inflammatory and HAND-associated pathways in both day-1 infected (HIV *pol* count > 5) and bystander microglia (HIV counts = 0), including interferon responses, TNFα/NF-κB signaling, IL-6/JAK/STAT3 signaling, and inflammatory response pathways (S7C Fig). Gene-level analysis of representative markers from these pathways further demonstrated overlapping expression distributions and comparable mean expression between infected and bystander cells at day 1 (S7D Fig). Furthermore, comparison of gene-level fold changes between bulk RNA-seq and scRNA-seq data showed strong agreement of changes in both day-1 infected (Spearman ρ = 0.91) and day-1 bystander (ρ = 0.86) populations (S7E Fig), indicating that the early inflammatory signature detected in bulk measurements at 24-hours post-infection reflects a population-wide microglial response rather than being driven exclusively by productively infected cells, suggesting an early sensing of the input virus by microglia.

### iMG exhibit a rapid and bi-modal inflammatory reaction following HIV infection

HIV infection of microglia may result in inflammatory responses that ultimately are damaging to surrounding glia and neurons and contribute to the development of HAND [44, 45]. We therefore sought to better characterize the principal inflammatory pathways that are activated over the course of HIV infection in iMG. Four major pathways as defined in Cluster 3 of Fig 3A were chosen: Inflammatory Response, TNFα signaling via NF-κB, IFNα response, and IL6/JAK/STAT3 signaling. Visualization of the temporal patterns shown as average FCs of genes within each pathway in iMG and MDMs clearly demonstrated the early (day 1) response across all pathways in iMG that was lacking in MDMs (Fig 4A). Following the early response, there was a decline in expression of inflammatory genes on day 2-4, followed by a gradual increase that was seen on day 6 and 8 to levels exceeding the day 1 response (Fig 4A). In infected MDMs, upregulation of inflammatory pathways was seen only on day 6 or 8, and the magnitude and breadth of the change in expression remained well below that seen in iMG (grey curves, Fig 4A).

**Fig 4.**
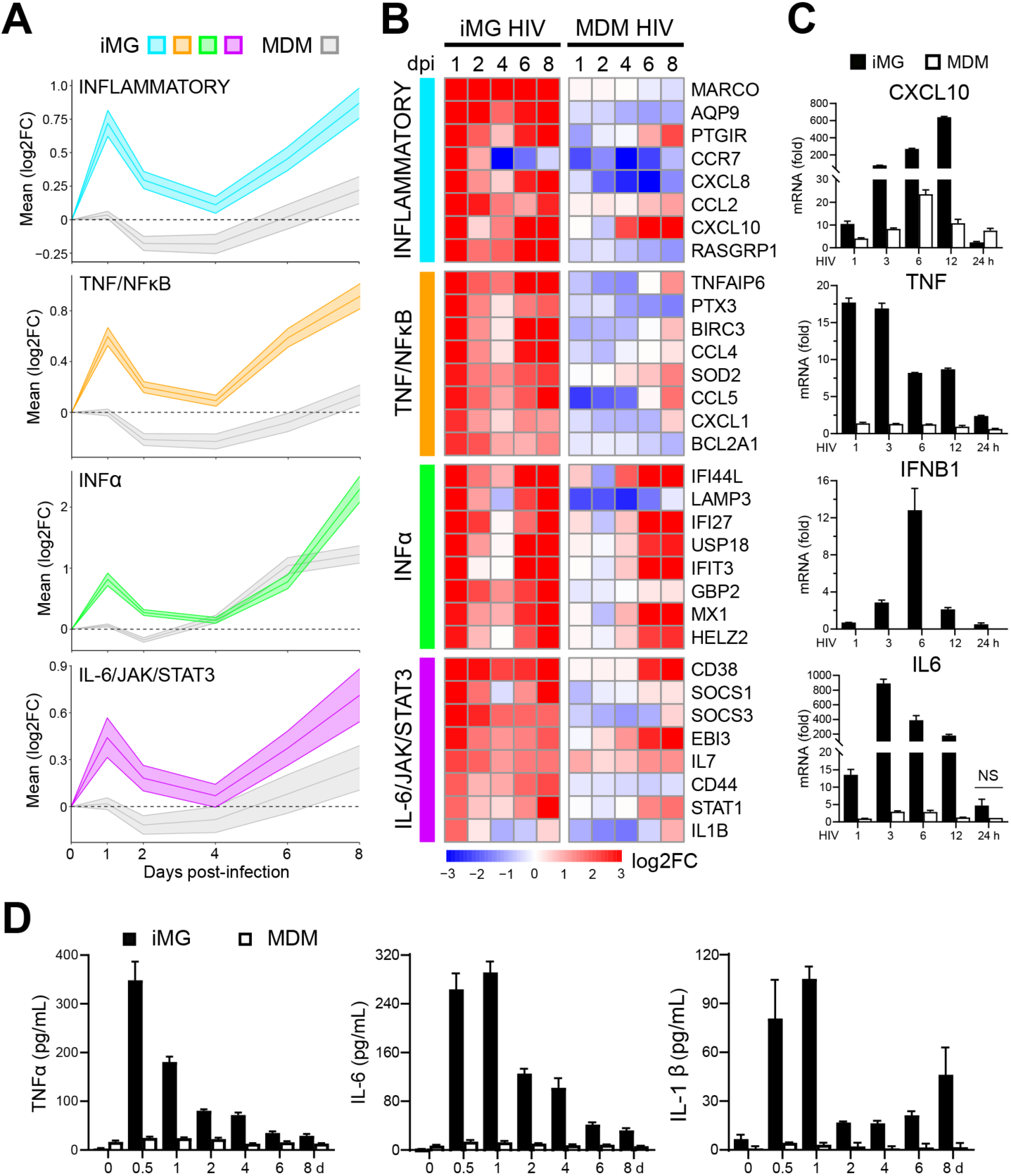
HIV-infected iMG develop rapid hyper-inflammatory responses. **(A)** Line graphs of average log_2_FC of all genes present in each inflammation-related pathway across 8-day HIV-infection time course in iMG. Shaded area represents the 95% confidence interval. Responses in MDMs are shown in grey. **(B)** Heatmap of temporal changes within iMG and MDMs in genes among selected inflammation-related Hallmark gene sets significantly enriched post-HIV infection as presented previously (Fig 3A). Top 8 genes with the greatest magnitude of expression changes on day 1 in iMG are represented from each pathway. **(C)** Bar graphs of gene expression changes for representative genes from each inflammation-related pathway quantified by RT-qPCR for iMG (solid black bars) and MDMs (white, black lined bars) at 1, 3, 6, 12 and 24-hours post-infection with HIV BaL. Statistical significance was determined using two-tailed, standard, unpaired t-test. Non-significance between iMG and MDM at a given timepoint post HIV-infection marked as NS, all other time points were significant (p≤0.05). **(D)** Bar graphs of cell culture supernatant cytokine levels measured for iMG and MDMs at 0.5, 1, 2, 4, 6 and 8-days post-infection with HIV BaL using Luminex-based multiplex assay. Statistical significance determined using two-tailed, standard, unpaired t-test and all differences shown between iMG and MDM were significant (p≤0.05).

Fig 4B depicts the most highly upregulated genes seen on day 1 post-infection from each of these pathways and demonstrates their change in expression over time following HIV infection using heatmaps. Elevation of TNF is a prominent feature of neuroinflammation seen in brain tissues from persons with HAND or HIVE [46]. NFκB gene expression itself was elevated in HIV-infected iMG, along with TNF receptor superfamily 9 message and the TNF-induced gene TNFAIP6 (Fig 4B, TNF/NFκB). Elevation of IFNα-induced genes including OASL, MX1, and IFI44L (Fig 4B, IFNα) is consistent with findings from brain tissue from persons with HIV and HAND or HIV, and with elevation of IFNα levels found in SIV-infected brain tissues [47–50]. The IL-6/JAK/STAT3 signaling pathway (Fig 4A-B, bottom panel) is a key driver of inflammatory and immune responses. In addition to the known role of STAT1 as already described, we note here that IL7 and IL1β gene products have been associated with neurological impairment in patients with HIV [46, 51], and CD38 is frequently seen to be elevated in neuroinflammatory and neurodegenerative conditions although not shown to be causal [52]. We conclude that the gene clusters shown in Fig 4A and 4B from HIV-infected iMG recapitulate many findings seen in brain tissues from individuals with HIV-associated neurocognitive disorders. To confirm the reproducibility of the biphasic inflammatory response observed in iMG following HIV infection, we conducted parallel experiments using a second, independently derived iPSC-microglia line (iMG2). The inflammatory gene expression dynamics in iMG2 closely paralleled those seen in the initial iMG line. Specifically, pathway-level analysis in iMG2 revealed early activation (day 1 post-infection) across the Inflammatory Response, TNFα/NFκB signaling, IFNα response, and IL-6/JAK/STAT3 pathways, followed by a transient decline (day 2), and a subsequent reactivation by days 6–8 (S8A Fig). These kinetics recapitulate the biphasic profile seen in the original iMG line and remain distinct from the delayed, attenuated response seen in monocyte-derived macrophages (MDMs). Gene-level expression profiles in iMG2 further validated these findings (S8B Fig). Characteristic early response genes such as MARCO, CXCL8, TNFAIP6, MX1, IFIT3, STAT1, IL1β, and CD38 all demonstrated comparable temporal trajectories to those observed in iMG (Fig 4B). The reproducibility of these patterns across two genetically distinct iMG lines strongly supports the notion that HIV elicits a robust, biphasic inflammatory program in microglia. This validation reinforces the utility of iPSC-derived microglia as a reliable in vitro model to dissect HIV-induced neuroinflammation.

To assess whether iMG exhibit a primed inflammatory phenotype prior to HIV infection, we compared the baseline (day 0) expression of inflammatory mediators across iMG, adult microglia (AMG), and monocyte-derived macrophages (MDMs) (S9A Fig). Genes were categorized into three functional groups: cytokines and receptors (CYT./RECEPTORS), sensory signal transducers (SENSORY TRANS.), and immediate/early cytokines (I.E. CYT). Across all three gene sets, we observed remarkably similar baseline expression patterns in iMG, AMG, and MDMs, suggesting that these cell types share a largely comparable resting inflammatory profile. While minor variations were observed, there was no evidence of elevated baseline activation in iMG relative to either primary adult microglia or monocyte-derived macrophages matured in GM-CSF or M-CSF. In contrast, LPS-stimulated MDMs (MDM2/M1, S9A Fig) exhibited the expected strong upregulation of inflammatory genes, validating the responsiveness of the markers analyzed. These data indicate that iMG do not exhibit an artificially heightened inflammatory state at rest and are instead transcriptionally aligned with primary, human adult microglia and macrophage populations in their basal inflammatory state. This supports the interpretation that the heightened responses observed after HIV exposure in iMG are not simply due to a pre-activated phenotype but reflect true, infection-induced inflammatory activation.

To further explore whether the heightened inflammatory response in iMG upon HIV infection reflects true transcriptional induction rather than differences in baseline gene expression, we stratified inflammation-related genes based on their relative expression at day 0 in iMG versus MDM. We then assessed their transcriptional trajectories over the course of an 8-day HIV infection using RNA-seq data. Genes that were more highly expressed at baseline in iMG relative to MDM and that fell within four inflammation-associated Hallmark pathways, Inflammatory Response, TNFα/NFκB signaling, IFNα response, and IL-6/JAK/STAT3 signaling were tracked over time (n = 421 genes). As shown in S9B Fig, these genes exhibited a biphasic induction pattern in iMG, with an early transcriptional spike on day 1 post-infection, followed by a partial decline on days 2 and 4, and a secondary increase by days 6 and 8. In contrast, MDMs showed minimal or delayed upregulation of these same genes, with modest increases only at later time points. Importantly, this pattern indicates that the dynamic upregulation in iMG extends well beyond what would be expected from modest baseline differences. Conversely, we analyzed the subset of genes that were more highly expressed at baseline in MDM relative to iMG, also within the same inflammatory pathways. As shown in S9C Fig, genes with higher basal expression in MDMs showed a similar expression pattern post-HIV infection in iMG. Together, these analyses demonstrate that the acute and sustained inflammatory activation observed in iMG following HIV exposure is not attributable to a pre-activated basal state, but rather reflects a cell-intrinsic, infection-specific response. The findings reinforce the distinction in inflammatory reactivity between iMG and MDM and highlight the unique sensitivity of iMG to HIV-induced neuroinflammatory triggers.

To validate the inflammatory signature detected via RNAseq and further delineate the kinetics of the rapid response observed in iMGs, we selected a single representative gene from each cluster to evaluate by RT-PCR (shown in Fig 4C). Analysis here was performed at multiple times during day 1 post-infection and compared to day 0 (uninfected) RNA levels. From the inflammatory cluster, we found that CXCL10 was rapidly induced in iMG, peaking at 12 hours post-infection, and showed significantly higher expression relative to infected MDMs. TNF RNA levels were very rapidly induced, as soon as 1-hour post-infection in iMG, while minor change was observed in MDMs. IFNB1 gene expression (representing type 1 IFN) rose and peaked at 6 hours in iMGs, and IL6 expression peaked at 3 hours post-infection (Fig 4C). Together, these results verify the difference seen in RNAseq studies in early responses between iMG and MDM, suggesting that there is a very rapid response in iMGs that is not seen in MDMs. Additional validation of the differences at early times post-HIV infection was then performed by measuring cytokine release by ELISA. Production of TNFα was rapid and robust, peaking at 12-24 hours post-infection in iMG (Fig 4D, left). Protein levels remained higher than those seen in infected MDM over the 8-day sampling period. IL-6 secretion was similar, peaking in the first day and remaining elevated as compared with MDMs throughout the time course (Fig 4D, middle). IL-1β levels in the supernatant peaked on day 1, then were lower on days 2-4 before rising again on days 6 and 8 (Fig 4D, right). Thus, RT-PCR and ELISA results confirm the early inflammatory response seen from RNAseq analysis in iMG that was lacking in HIV-infected MDM.

### Receptor and restriction factor expression is rapidly induced in iMG upon HIV infection

Although our primary focus in this study was on the genes and pathways in HIV-infected microglia that are likely to provide insights into HAND, we sought also to define the expression of a panel of HIV-relevant genes following infection of iMG. We first analyzed basal expression of receptor CD4, coreceptors CCR5 and CXCR4 and a panel of restriction factor genes in iMG and MDMs (Fig 5A). Receptor and coreceptor messages were significant at baseline in both MDMs and iMG, with reduced levels of CXCR4 in iMG as compared with MDMs and compared with the published data for AMG (Fig 5A). Expression of CD4 and CCR5 in all cell types was consistent with prior results indicating that receptor and coreceptor levels do not limit infection of microglia with M-tropic, R5 HIV [29]. Upon HIV infection, HIV receptor and coreceptor genes increased in RNA expression from day 1 to day 8 in iMG and MDMs (Fig 5B). Siglec-1 has been shown to play a key role in HIV transmission through trans-infection from both DCs and macrophages to T cells, and contributes to the formation of the virus-containing compartment (VCC) in macrophages [53–55]. Interestingly, SIGLEC1 expression levels were elevated on day 1 in iMGs (FC of 2.0) but less so in MDMs. SIGLEC1 expression continued to rise in iMG to a log_2_FC of 4.1 on day 8 (Fig 5B).

**Fig 5.**
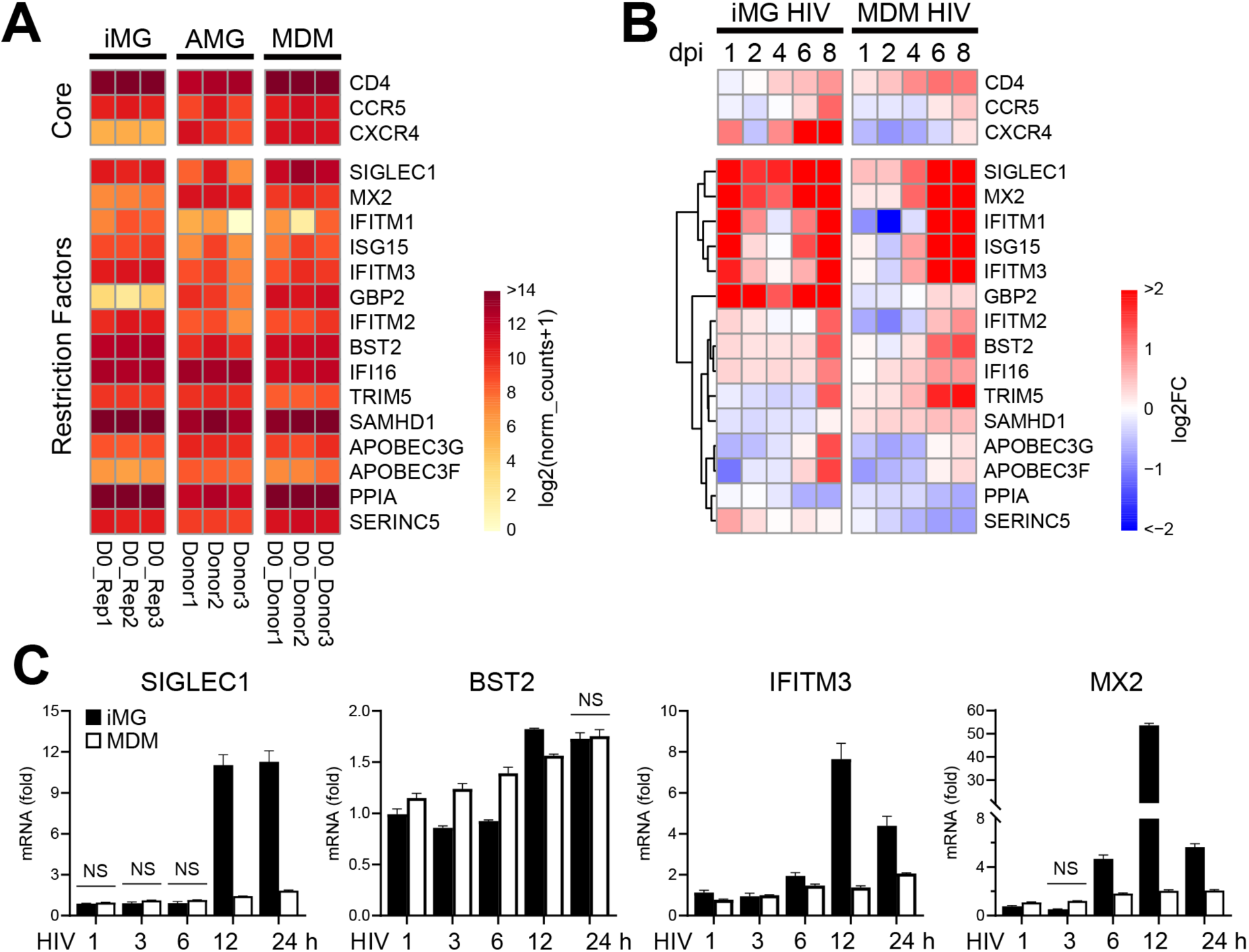
Infection of iMG induces rapid gene expression changes in HIV-related factors. **(A)** Steady state (day 0) gene expression levels for a panel of HIV-related factors in iMG (n = 3 biological replicates), adult microglia (AMG) and MDMs (n = 3 donors). Color in heatmap represents log_2_ (normalized counts+1). **(B)** Gene expression changes in HIV-related factors across 8-day time course of infection at an MOI of 0.25 in iMG and MDMs. Color represents log_2_ fold change. **(C)** RT-qPCR quantitation of mRNA fold changes at 1, 3-, 6-, 12- and 24 hours post-infection for SIGLEC1, BST2, MX2 and IFITM3 (technical replicate shown, 2 biological replicates performed with no magnitude or temporal differences). Statistical significance was determined using two-tailed, standard, unpaired t-test. Non-significant differences between iMG and MDM at a given timepoint post HIV-infection are marked as NS, all other time points demonstrated significant differences (p≤0.05).

Restriction factors are innate cellular defense proteins that inhibit the replication of viruses for the benefit of the host [56, 57]. We analyzed gene expression changes in a panel of myeloid-relevant restriction factors following infection of iMG and MDMs (Fig 5A and B). HIV infection caused a rapid induction of restriction factor gene expression in iMG on day 1, with six restriction factors exhibiting a log_2_FC greater than 1.5 (Fig 5B). Restriction factors exhibiting a log_2_FC of greater than 3 in infected iMG included MX2, IFITM1, GBP2, AND ISG15. In contrast, no restriction factors in infected MDMs were induced above a log_2_FC of 0.5 on day 1 or day 2. Expression of most restriction factors increased more gradually in MDMs, so that both iMG and MDMs demonstrated increases on day 6 and day 8 (Fig 5B). One of the differences observed between HIV-infected iMG and MDMs was seen with the IFN-inducible factor GBP2, encoding a protein that restricts Env protein processing and incorporation [58], which was expressed at low levels in iMG at baseline but was rapidly upregulated following infection. It is also notable that SAMHD1, a dNTPase that limits HIV-1 infection of macrophages and resting T cells [59, 60], was highly expressed at baseline but rose following infection only in MDMs (Fig 5B). BST2 encodes the restriction factor tetherin, a type 2 transmembrane protein that imposes a late restriction on particle release [61]. BST2 was well expressed at baseline and upregulated at late time points following infection in both iMG and MDM (Fig 5B). These data indicate that HIV infection of iMG induces rapid expression of multiple restriction factors, whereas comparative induction in MDM is more delayed. The early induction of a subset of IFN-stimulated restriction factor genes in iMG was confirmed by RT-PCR (Fig 5C). Increases in BST2 expression were comparable in both cell types, while early increases in iMG were confirmed for SIGLEC1, IFITM3, and MX2 and were lacking in MDMs (Fig 5C). Early induction of restriction factors may be important in some contexts, although in these *in vitro* experiments it did not appear to limit the magnitude of infection when compared to that of MDMs (Fig 2).

### HIV infection induces gene expression changes in iMG that recreate HAND signatures

HIV-infected microglia have been implicated in the pathogenesis of HAND. To investigate the relationship between the observed HIV-induced inflammatory responses in iMG and established signatures of HAND, we performed GSEA on the iMG transcriptional response using previously identified HAND genetic signatures defined from postmortem brains of PLWH showing HAND symptoms [62, 63] and CNS tissues of SIV-infected non-human primates [64], along with modules derived from the MGEnrichment application [65]. Modules containing genes upregulated in postmortem brain tissue of patients with HAND and HIV encephalitis (HIVE), as well as from brains of SIV-infected non-human primates, were consistently positively enriched in iMG following HIV infection *in vitro* (Fig 6A, HAND cluster). The upregulation of genes associated with HAND was apparent at the earliest timepoint following HIV infection of iMG (day 1) and was also present at late time points. Some of the early responses in infected iMG were also positively enriched in published signatures from HIV-uninfected brains that are associated with neurocognitive disorders (Fig 6A, neurocognitive cluster), including Alzheimer’s disease (AD>Control) and inflammaging (Aging_inflammatory) [66, 67]. We then selected ‘core’ HAND signature genes that were shared by 2 or more HAND modules (a total of 79 genes) and analyzed their expression in HIV-infected iMG across the time course. This analysis provides a clear view of the peak in expression of HAND-related genes during the early (day 1) response, followed by a decline and then a gradual rise from day 4 to day 8 (Fig 6B). Shown in red are the ten most highly induced genes on day one from infected iMG. These represent key genes that have previously been indicated as biomarkers of HAND and/or HIVE, including IFI44L, IFI44, ISG15, STAT1, MX1, IFIT3, PARP9 and NLRC5, along with the IFN-induced HIV-1 attachment/transmission factor SIGLEC1, and are all type I ISGs.

**Fig 6.**
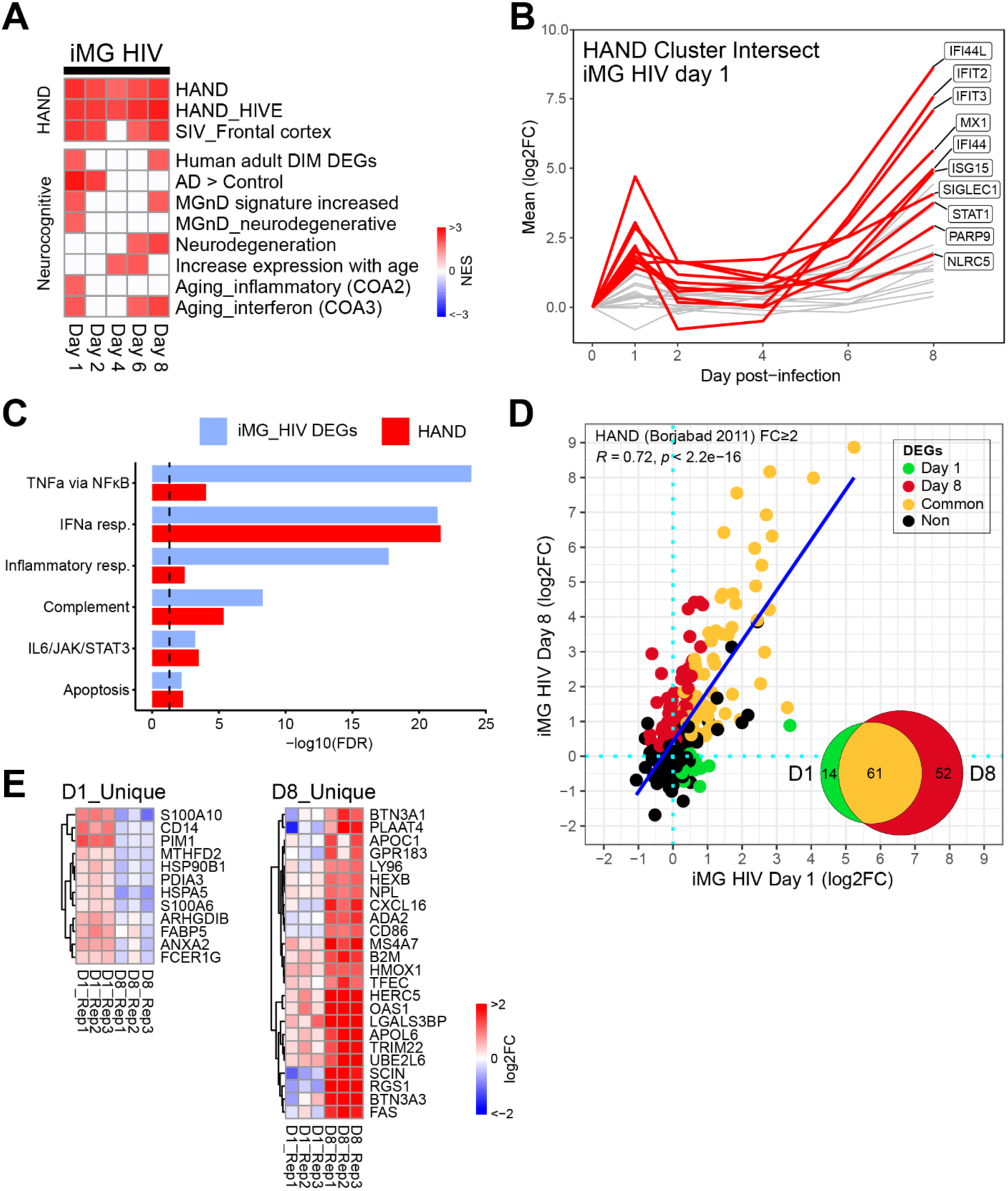
Genes linked to HAND are upregulated early and late in HIV-infected iMG. **(A)** Analysis of gene set enrichment in HIV-infected iMG across 8-day time course using curated microglia disease state modules from the MGEnrichment application [65] and HAND-associated modules [62, 63]. Modules are listed in S4 Table. Color represents the NES of significantly enriched modules (FDR<0.05, absolute NES>2). **(B)** Line graph depicting the average fold changes of 79 genes found in two or more (≥2) HAND modules from HIV-infected iMG. The genes highlighted in red represent the top 10 genes included in two or more of the HAND modules on day one post-infection. **(C)** ORA comparing Hallmark pathway enrichment of HAND genes from Borjabad 2011 et al. [62] and DEGs from HIV-infected iMG. Bars depict the FDR significance levels, while the red dashed line denotes the threshold for significance (FDR<0.05). **(D)** Scatter plot of fold changes in upregulated HAND genes from [62] that were also identified on days 1 (x-axis) and 8 (y-axis) post-HIV infection in iMG. HAND-associated genes identified in iMG were colored according to time post-infection at which they were differentially expressed (red = day 8 unique, green = day 1 unique, yellow = common). HAND-related genes were identified as DEGs in our study using FDR < 0.1 criteria. Venn diagram in bottom right corner displays quantification of unique DEGs to either day 1 or day 8 and common DEGs (48% common to both day 1 and day 8 post-infection). **(E)** Distinct expression changes of HAND genes from Borjabad 2011 following HIV infection of iMG. The heatmap illustrates genes that were uniquely upregulated on either day 1 (D1_Unique) or day 8 (D8_Unique) post-infection (n=3 biological replicates). Color represents log_2_ fold change.

Next, we performed Hallmark pathway enrichment of the Borjabad HAND signature gene set [62] and the HIV-infected iMG DEGs from the current study. This analysis revealed multiple co-enriched pathways for HAND and HIV-infected iMG, including interferon alpha responses, complement activation, TNFα / NF-κB signaling, IL6/JAK/STAT3 signaling, hypoxia, inflammatory responses and apoptosis (Fig 6C). The striking similarity of signatures between infected iMG and the HAND signature was further assessed by intersecting the genes that were differentially expressed in both datasets. There was significant overlap between the DEGs in the two conditions, with 57% (138/243 genes) of the postmortem HAND signature also induced by HIV infection of iMG, and the overlap increased over time (S10 Fig). We then examined those shared upregulated HAND signature genes present on day 1 post-infection with those expressed later in infection (day 8). A comparison of these shared HAND genes revealed a strong correlation between the early and late expression of this inflammatory signature (R=0.72, Fig 6D). Among the HAND genes upregulated in HIV infected iMG on day 1 or day 8, the greatest portion (61/127) were common to both timepoints. The initial response displayed a small subset (14/127) of uniquely induced HAND genes, while the later response had a greater unique response (52/127) (Fig 6D, Venn diagram). Visualization of the expression of these common HAND-associated genes confirmed that while many HAND signature genes were induced in both stages of the response, distinct subsets were present in iMG exclusively at early vs. late times post-HIV infection (green vs red, Fig 6D). Unique genes for day 1 and day 8 are shown in Fig 6E. The early response unique HAND DEGs included CD14, S100 proteins (S100A10 and S100A6), and PIM-1, a kinase involved in HIV-1 latency [68]. Late unique response genes included those encoding the inflammatory chemokine CXCL16; the cell surface death receptor gene FAS; OAS1, a gene often implicated in HAND pathogenesis [50]; TRIM22, an ISG that can inhibit HIV transcription and promote latency [69]; and HERC5, encoding an E3 ligase linked to inhibition of the late stages of HIV replication [70]. Common genes to the two time points included numerous ISGs, including ISG15, CCL8, IFI44L, MX2, IFITM1, GBP1, GBP2, and SIGLEC1, in addition to STAT1 and TNF superfamily member TNFSF13B. Together, these results demonstrate that HIV infection of iMG *in vitro* was able to recapitulate, in large part, the HAND immune signature derived from postmortem human brain.

## Discussion

The pathogenesis of HAND remains incompletely understood. HIV can be detected in the CNS along with elevated inflammatory markers as early as 8 days following infection [71]. The initial entry of HIV into the CNS is likely to occur through transit of infected T cells and monocytes, followed by infection of brain resident cells such as perivascular macrophages or microglia [72–74]. Breakdown of the blood-brain barrier through the action of viral proteins has also been suggested to enhance HIV entry into the CNS [75–77], while alternatively transcellular migration of infected CD4+ T lymphocytes across an intact BBB has been reported to contribute to HIV neuropathogenesis [78]. Inflammation in the CNS is triggered soon after entry of HIV, and the detection of elevated inflammatory markers in CSF and in postmortem brain tissue is a hallmark feature of HAND [46, 51, 71]. Microglia are the major resident myeloid cells of the brain and are highly susceptible to HIV infection. Infection of microglia has been detected at early time points following infection of rhesus macaques with SIV and resulted in inflammatory or damage-associated microglial phenotypes [79, 80]. Microglia have been demonstrated to be infected in human brain even at early times post-infection, harbor replication-competent virus, and are thought to contribute to the latent viral reservoir [6]. Given the prominent role of microglia in early and chronic HIV infection and their potential to elicit the inflammatory responses that contribute to neuronal damage leading to HAND, we designed this study as a comprehensive analysis of the gene expression changes that occur in microglia following HIV infection, using an iPSC-derived microglial model system that has been previously established to closely represent authentic human microglia [27–29]. Gene expression studies were conducted in parallel using HIV-infected human MDMs to distinguish the features of microglia infection that are held in common with macrophages, and to identify those responses that are unique to microglia.

A major finding from this study is the strong overlap of the inflammatory profiles seen in HIV-infected microglia following *in vitro* infection with an M-tropic HIV strain to those that have come from postmortem HIV-infected brain tissues derived from persons with HAND or HIVE. We note that HIVE is distinct from HAND, with HIVE now a rare neurologic manifestation in the post-cART era, while some of the datasets used here and elsewhere have overlapping inflammatory features. Beginning even within the first 24 hours of infection, iMG demonstrated upregulation of inflammatory genes in a signature that has been previously identified as characteristic of HAND. Single-cell transcriptomic analyses on day 1 post-infection revealed that the robust inflammatory response observed at day 1 post-infection was broadly induced across the microglial population, including both the minority of productively infected and bystander cells. The strong concordance between bulk and scRNA-seq profiles at this early time point indicates that HIV exposure rapidly elicits a coordinated, population-wide microglial inflammatory program that mirrors neuroinflammatory signatures associated with HAND. Following this early response, concordance with the HAND signature grew more pronounced as virus spread through the population, reflecting both a rising number of shared genes and their higher expression over time (Fig 6D, S10 Fig).

Prominent among the clusters of upregulated genes were those involved in IFN signaling, TNFα/NFκB signaling, and the IL-6/JAK/STAT3 pathway, as well as inflammatory chemokines including CXCL8 and CXCL10. While many of the individual genes upregulated in this study have been implicated as mediators of inflammation in HIV-infected brains or from CSF-based studies from PLWH with clinical findings of HAND, the significant overlap with the HAND signature profile is a new finding.

The pattern of gene expression following HIV infection of iMG was distinctly different than that of macrophages. Infection of iMG produced a very early upregulation of inflammatory pathways, suggesting an early signaling event that was absent in MDMs, and the magnitude (FC) of differential expression of inflammatory genes was significantly higher than that seen in MDM. The early activation followed by a decline and later rise in expression was found in common to several inflammatory pathways, including IFN-mediated signaling, TNF/NFκB signaling, and IL-6/Jak/Stat pathways. IFN-mediated signaling and TNFα elevation are hallmark findings in HAND. IFNα has been found to be elevated in the CSF of HIV-infected persons with dementia and has been suggested to have a role in the pathogenesis of HAD [81–84]. Elevated CSF IFNα has also been implicated in more subtle neurocognitive impairment in HIV-infected individuals [85]. Similarly, TNFα is a key mediator of inflammation that has been found to be consistently elevated in postmortem brain tissue from patients with HAND [46].

While the early and robust inflammatory profile and strong overlap with HAND/HIVE signatures following infection of iMG are new findings, previous infection studies performed *in vitro* in microglia have reported upregulation of some of the same inflammatory pathways we highlight in this work. Boreland and colleagues focused on the sustained type 1 IFN-mediated signaling that occurred following HIV infection of iMG [86]. These investigators reported the upregulation of ISGs IFIT3, MX2, ISG15, IRF7, OAS1, STAT1, CXCL10, IFITM3, and SIGLEC1, consistent with the work shown here. They also showed upregulation of CCL2, TNFα, and IL-1β, while not observing upregulation of IL6. Kong and colleagues examined HIV infection of microglia in a cerebral organoid model, and demonstrated enhanced production of CXCL10, CCL2, and a series of ISGs including MX1, ISG15, ISG20, IFI27, and IFITM3 [44]. In an iMG/neuron co-culture model, Akiyama and colleagues have shown that HIV infection upregulated CXCL10, CCL2, CCL7 and ISGs SIGLEC1 and RSAD2, and demonstrated that intron-containing RNA export into the cytosol was required for induction of proinflammatory responses [87]. In a humanized mouse model utilizing iMG and engrafted human PBMCs, Min and colleagues demonstrated elevation of TNFα, IL-6, and CD68 message following HIV infection [88]. A neuroinflammatory profile was also seen in a tri-culture method employing iMG with stem cell-derived neurons and astrocytes, including elevation of IL-1β, TNFα, and IL-8 [89]. Thus, there are clear and consistent results indicating that HIV infection of iMG stimulates inflammatory pathways, and that enhanced expression of ISGs and TNFα is a prominent feature. In the current study, we have presented a global picture of gene expression in HIV-1-infected iMG over a time course of acute infection, together with a myeloid comparator (MDM) to derive features unique to iMG. The extensive overlap at both early and later time points in HIV-infected iMG with gene expression studies from postmortem brain tissues of individuals with HAND or HIVE provides suggestive evidence that microglia represent the major source of neuroinflammation in humans with HAND, affirming the relevance of this *in vitro* infection model. This study highlights the importance of IFN-stimulated pathways as well as TNF/NFκB and IL-6/JAK/STAT signaling in microglia, demonstrating that these inflammatory signaling pathways are triggered earlier and to higher degree than that seen in HIV-infected macrophages. This supports the idea that microglia are uniquely susceptible to HIV-induced inflammatory responses, as compared to tissue macrophages in other compartments, and this may help explain the prominence of HAND in HIV-infected individuals even following institution of cART. A heightened susceptibility to inflammation may require only transient or low levels of viral replication to elicit, leading to ongoing neural damage over time.

Chronic inflammation is a major driver of the neuroaging phenotype, contributing to cognitive decline, synaptic dysfunction, and increased vulnerability to neurodegenerative diseases such as HAND and Alzheimer’s Disease. HIV infection of iMG showed significant enrichment for gene sets associated with both a neurodegenerative and senescent or aged phenotype [14, 30, 66, 90]. Targeting inflammatory pathways and modulating microglial function may therefore represent promising therapeutic approaches to mitigating the detrimental effects of inflammaging and preserving cognitive function in PLWH.

Data in this report also elucidates the expression of HIV-relevant genes in iMG and changes occurring following HIV infection. One of the interesting findings is that SIGLEC1 RNA expression is relatively high at baseline in iMG and then is rapidly increased following HIV infection. Expression of SIGLEC1 is not limited to the iMG system, as similar levels have been documented from brain-derived AMG [27]. The expression of Siglec-1 at the protein level has been reported to be very low or undetectable in microglia within mouse brain outside of the choroid plexus and leptomeninges but can be induced following damage to the blood-brain barrier [91]. SIGLEC1+ microglia have been seen in brains of humans with neuroinflammation from multiple sclerosis [92] and have been implicated in neuroinflammation in a murine model of ceroid lipofuscinosis [93]. The rapid upregulation of SIGLEC1 expression shown in the present study is of interest, not just as a potential marker of neuroinflammatory microglia phenotype, but also because this lectin plays a major role in HIV-myeloid cell interactions. In DCs and MDMs, Siglec-1 specifically captures HIV-1 particles through interactions with gangliosides incorporated onto the viral lipid envelope, leading to coalescence of Siglec-1-HIV microdomains on the cell surface and subsequent internalization of viral particles to a compartment known as the virus-containing compartment or VCC [53–55, 94]. The VCC functions as a holding compartment in these cell types and can contribute to trans-infection of susceptible cells upon cell-cell contact. The high expression of Siglec-1 in HIV-infected microglia might similarly be expected to facilitate viral capture and spread, potentially contributing to the spread of HIV infection within the CNS.

BST2/tetherin is an IFN-induced host restriction factor that arrests the final release of viral particles through the presence of two membrane anchors, a membrane-spanning domain and a GPI anchor, allowing attachment to both the viral lipid envelope and the plasma membrane [61]. The N-terminus of tetherin contains a hemiITAM motif that upon particle retention and dimerization of tetherin is phosphorylated, recruiting the spleen tyrosine kinase (Syk) and activating NF-κB signaling [95]. We found that tetherin/Bst2 is upregulated at late time points in HIV-infected microglia. Although the accessory protein Vpu of HIV downregulates tetherin and thus inhibits tetherin-mediated signaling, macrophage-tropic viral isolates including some derived from the CNS often are defective in *vpu* expression [96]. It will be interesting in future studies to decipher the contribution of tetherin-mediated signaling to neuroinflammation in HIV-infected microglia, as this may provide a direct link between active viral replication and signaling leading to activation of NF-κB.

Results here together with those from other groups highlight the high susceptibility of iMG to HIV-1 infection [29, 44, 86, 88, 89]. We found that viral infection and subsequent replication were robust in iMGs even at low MOI, using an R5 M-tropic HIV-1 isolate, indicating that receptor CD4 and coreceptor CCR5 were not limiting. Indeed, at the RNA level and at the surface protein level expression of CD4 and CCR5 are robust in iMG [29]. AMG from human brain also indicate strong expression of CD4 and CCR5 [27]. CD4 levels in microglia have been reported by some to be undetectable in fresh brain tissue from HIV-negative individuals, while detectable at low levels in microglia from HIV-infected individuals [97]. However, others have shown that *ex vivo* brain-derived microglia express surface CD4 at levels comparable to that of MDM, and shown that CD4 and CCR5 levels are greatly upregulated in response to inflammatory cytokines including IL-1β, IL-6, and TNFα [98]. Some of the discrepancy in reported CD4 levels in microglia may come from how *ex vivo* microglia are handled, with activation following isolation potentially leading to higher CD4 expression. It is reasonable to postulate that initial infection of a limited number of microglia may lead to inflammatory responses that result in upregulation of CD4 in surrounding cells and thereby enhance susceptibility to the spread of infection.

We found a robust induction of inflammatory signatures in iMG following acute HIV infection that differed from that in MDM. The sensor(s) responsible for the initial rapid induction of innate responses in iMG remain under evaluation. Innate immune sensing of HIV-1 in its target cells can occur through a variety of mechanisms. TGFβ-activated kinase (TAK1) activation can occur at early times following viral entry through TRIM5 interactions with the incoming capsid [99] or potentially through interaction of the cytoplasmic domain of gp41 [100]. RIG-I-dependent sensing of HIV-1 genomic RNA secondary structures in the cytosol of infected T cells and macrophages has been reported, leading to robust expression of ISGs [101, 102]. cGAS has been shown to sense HIV cDNA in CD4+ T cells and myeloid DCs [103–105]. Plasmacytoid DCs initiate an early type I interferon response stimulated by incoming HIV-1 single-stranded RNA by TLR7 [106, 107]. Importantly, McCauley and colleagues reported that viral integration followed by production and nuclear export of intron-containing RNA was required for activation of IFN and inflammatory pathways in T cells and macrophages [108]; this work was followed by evidence that MDA5 was required for sensing of the intron-containing RNA in myeloid DCs and macrophages [109]. Akiyama and coworkers reported that nuclear export of intron-containing RNA was required for induction of inflammatory responses in iMG, establishing this mechanism as an important one in this cell type [87]. The early inflammatory responses observed in our study, together with the data cited above suggest that there may be at least two sensing events in iMG from the present study, one during early stages following virus entry that results in an early (day 1) peak response, followed by a slower rise over the course of HIV replication that is most likely explained by production of intron-containing HIV RNA and sensing through MDA5 as previously reported. However, the sensors responsible for the kinetics of inflammatory responses observed here will need to be directly addressed in future studies.

One of the previous differences noted in HIV infection of microglia vs. MDMs is the higher susceptibility of microglia to HIV-1 infection and spread as compared to macrophages [110]. This high susceptibility has been linked to differences in SAMHD1-mediated restriction. SAMHD1 exhibits higher levels of phosphorylation in microglia, inhibiting its dNTPase activity and limiting restriction of HIV-1 [87]. Inflammatory responses mediated by icRNA were found in this study on day 6 following infection of iMG with HIV. Some of the differences in inflammatory profiles observed in spreading culture at later timepoints in our study could be influenced by this differential ability to infect and spread in microglia vs. MDMs, although we did not see significant differences at most time points in viral replication as indicated by p24 release. The early timepoint peak of inflammation (day 1) seen in the present study, however, would not likely have been a result of icRNA production, but is more likely due to sensing of incoming virions. Another important consideration for the interpretation of the present study is that co-culture systems may result in a more homeostatic state of microglia [111]. Assessment of the potential modulating effects of co-culture with neurons as well as other mixed culture systems on inflammatory pathway activation in microglia following HIV infection will be an important to establish in future studies.

It will be of great interest to define how HIV alters transcriptional regulatory networks in infected microglia in future studies. The epigenetic changes induced by HIV following HIV sensing may lead to sustained changes in a subset of microglia that can contribute to neuroinflammation and neuronal damage during chronic infection or following institution of cART and suppression of viral replication. Inflammation may alter chromatin accessibility of regulatory regions of genes involved in inflammation, potentially leading to a sustained proinflammatory phenotype or a tendency to develop an enhanced response upon restimulation [112]. Understanding epigenetic changes will require combining comprehensive gene expression analysis with studies of chromatin accessibility and transcription factor binding in relevant infection models. While isolated microglia or iMG can reveal key aspects of the changes occurring following HIV infection, it will also be useful to perform this analysis within the context of 3D organoid models and relevant animal models, and to do so in the context of chronic infection and antiretroviral therapy.

## Materials and Methods

### Induced Pluripotent Stem Cell (iPSC) Culture and Maintenance

iPSC_TFS (hPSCreg: TMOi001-A, female, used to derive iMG1 cells shown in all figures) and iPSC_72.3 (hPSCreg: CCHMCi001-A, male, used to derive iMG2 cells were employed in this study (S5 Fig and S8 Fig). iPSC_72.3 was derived from primary human foreskin fibroblasts (HFFs) cultured from neonatal human foreskin tissue. Tissues were obtained through the Department of Dermatology, University of Cincinnati. iPSC_72.3 was generated by the CCHMC Pluripotent Stem Cell Facility, approved by the CCHMC institutional review board, and previously characterized [113, 114]. iPSC_72.3 demonstrated normal karyotype and differentiated into endoderm, mesoderm, and ectoderm lineages in an *in vivo* teratoma assay. iPSC TFS was derived from umbilical cord blood, obtained through Thermo Fisher and displayed a normal karyotype. iPSCs were cultured on Cultrex (R&D Systems) coated plasticware and maintained in mTeSR1 (STEM Cell Technologies) at 37 °C with 5% CO2. Cells were passaged using Gentle Cell Dissociation Reagent (STEM Cell Technologies) according to manufacturer’s instruction. Karyotype and mycoplasma contamination were routinely monitored.

### Production of Hematopoietic Progenitor Cells (HPCs)

Differentiation of iPSCs to HPCs was performed using the STEMdiff Hematopoietic Kit (STEM Cell Technologies). On the day prior to differentiation, iPSCs were passaged with Gentle Cell Dissociation Reagent (STEM Cell Technologies) and aggregates were plated onto several 10 cm^2^ dishes at a target density range of 5 – 10 aggregates per cm^2^. On the following day, plates containing between 80 and 100 total colonies (approximately 2 per cm^2^) were chosen to proceed with differentiation protocol. On day 0, mTeSR1 media supplemented with 0.5 µM Thiazovivin was replaced with 8 mL of basal media A containing a 1:200 dilution of supplement A. On day 2, a half culture basal media A change was performed. On day 3, basal media A was completely removed and replaced with basal media B containing a 1:200 dilution of supplement B. Half media exchanges were performed with basal media B on days 5, 7, 9, 10, 12 and 14. On days 12, 14 and 16 non-adherent cells were collected and centrifuged for 5 min at 300 x g. Clarified supernatants were half media exchanged with basal media B on days 12 and 14 after harvesting HPCs. Pelleted cells were cryopreserved at a density of 1.5 – 2 x 10^6^ HPCs per mL of CryoSTOR CS10 (STEM Cell Technologies).

### iPSC-derived microglia (iMG) Maturation and Culture

Frozen HPCs were thawed rapidly by immersion in a 37 °C water bath and cultured immediately in cytokine supplemented iMG maturation media and plated onto 1 mg/mL growth factor reduced (GFR) Matrigel-coated 6-well plates at 1.5 x 10^5^ cells per cm^2^ or 2.5 x 10^5^ per well. HPCs are cultured at a density of 1.5 × 10^5^ per cm^2^ onto 1 mg/mL GFR Matrigel-coated 6-well plates in 2 mL of iMG media per well (DMEM/F12, 2% B27, 0.5% N2, 2% insulin-transferrin-selenium, 1X MEM Non-Essential Amino Acids Solution, 1X GlutaMAX, 400 µM 1-thioglycerol and 5 µg/mL human insulin). Prior to use, iMG media was supplemented with 100 ng/mL human IL-34, 50 ng/mL TGF-β1 and 25 ng/mL M-CSF (Peprotech). On days 2, 4 and 6 media were supplemented with the addition of 1 mL per well of iMG media with cytokines. On day 8, media was removed leaving behind 1 mL per well of conditioned media. Cells were centrifuged for 5 min at 300×g, media aspirated, and cells resuspended in 1 mL of iMG media with cytokines prior to addition back to wells. On days 10, 12 and 14 media were again supplemented with addition of 1 mL per well of iMG media with cytokines. On day 16, media was removed leaving behind 1 mL per well of conditioned media. Cells were centrifuged for 5 min at 300×g, media aspirated, and cells resuspended in 1 mL of iMG media with cytokines prior to addition back to wells. On days 18, 20 and 22 media were again supplemented with addition of 1 mL per well of iMG media with cytokines. On day 24, cells were resuspended in iMG media supplemented with a five-cytokine cocktail consisting of 100 ng/mL human IL-34, 50 ng/mL TGF-β1, 25 ng/mL M-CSF, 100 ng/mL CD200 and 100 ng/mL CX3CL1 to facilitate final maturation into iMG. On days 26 and 28, cells were fed by the addition of 1 mL per well of iMG media supplemented with a five-cytokine cocktail. By day 28 iMG were considered mature and used for further characterization and RNA seq analyses. Cells were maintained for a maximum of 2 weeks following the 28-day cycle of maturation and conditioned media maintenance described above. Reagents and media formulations used to generate iMG are listed in S1 Table.

### Human Whole Blood-derived Monocyte Isolation and Monocyte-derived Macrophage (MDM) Maturation and Culture

Human peripheral blood mononuclear cells (PBMCs) were isolated from fresh heparinized blood by Ficoll-Hypaque gradient centrifugation. Buffy coats were pooled, and platelets removed by washing repeatedly with phosphate-buffered saline (PBS). Monocyte enrichment was performed by indirect magnetic labeling using the Pan Monocyte Isolation Kit (Miltenyi Biotec) according to manufacturer’s protocol. Enriched monocytes were plated on poly-D-lysine coated plates (Corning) and type 1 rat tail collagen coated 35 mm MatTek dishes (MatTek). Monocytes were maintained in RPMI-1640 supplemented with 10% FBS (Lot No., R&D Systems), 100 µg/mL streptomycin, 100 U/mL penicillin, 2 mM GlutaMAX, and 5 ng/mL GM-CSF (Peprotech). Monocyte cultures were maintained in GM-CSF supplemented media for 7 days to mature cells to MDMs. Media was replaced every 3–4 days.

### Production of Cell-free Stocks of HIV-1 BaL in PBMC and Infection

HIV-1 isolate BaL stocks were prepared as follows: Human peripheral blood mononuclear cells (PBMCs) were isolated from fresh heparinized blood by standard Ficoll-Hypaque gradient centrifugation methods. PBMCs were resuspended in RPMI 1640 supplemented with 20% heat-inactivated fetal bovine serum and 50 μg/mL gentamicin (RPMI 1640-GM). Primary HIV-1 isolates were propagated in PBMCs stimulated with 5 μg/mL phytohemagglutinin (PHA-P, Sigma-Aldrich) and 5% IL-2/T-cell Growth Factor (Hemagen Diagnostics). The IL-2/PHA-stimulated cells were infected using a high-titer seed stock of virus minimally passaged in PBMCs, starting from a viral stock obtained through the NIH AIDS Reagent Program (from Dr. Suzanne Gartner, Dr. Mikulas Popovic and Dr. Robert Gallo). One mL of virus was transferred to the flask containing freshly stimulated PBMCs and incubated overnight at 37 °C in 5% CO2. The cells were washed and resuspended in 30 mL of RPMI-GM supplemented with 5% IL-2/T-cell Growth Factor (Hemagen Diagnostics). Typically, the virus was harvested two times; the first harvest was on day 4 post-infection, with subsequent harvest on day 7. The virus-containing supernatants were collected, clarified by centrifugation, and filtered through a 0.45-μm filter. The virus was then aliquoted into 2 mL sterile screwcap cryovials and stored at − 80 °C. iMGs and MDMs were infected with primary HIV-1 isolate BaL at an MOI of 0.05, 0.1, 0.25 and 0.5 for generating multiple round growth curves over 14 days and an MOI of 0.25 for experiments involving subsequent RNA seq analyses.

### Titration of HIV-1 in TZM-bl Cells

Infectivity of viral stocks were assayed for infectivity using TZM-bl indicator cells (obtained through the NIH AIDS Reagent Program, Division of AIDS, NIAID, NIH; from Dr. John C. Kappes, Dr. Xiaoyun Wu and Tranzyme Inc.). TZM-bl were incubated for 48 h with a serial dilution of HIV-1 BaL stocks, and 100 μL of supernatant was removed from each well prior to the addition of 100 μL of Britelite Plus substrate (Revvity). Measurement of infectivity involved transfer of 150 μL of cell/substrate mixture to black 96-well solid plates and measurement of luminescence.

### P24 ELISA

P24 antigen content of infected cell supernatants were measured using a p24 antigen capture ELISA as described previously [53]. Briefly, murine anti-p24 CA183 hybridoma supernatants were coated onto MaxiSorp 96-well plates (Nunc) at a dilution of 1:800 in PBS and incubated overnight at 37 C. Plates were washed two times with PBS and blocked for 1 h at 37C with 5% fetal calf serum (FCS) in PBS. Samples were diluted in p24 ELISA sample diluent containing 5% FCS and 0.5% Triton X-100 in PBS and incubated for 2 h at 37 C. Plates were then washed four times with 0.1% Tween 20 in PBS. The detection of bound p24 was determined using HIV-Ig, obtained from NABI through the NIH AIDS Research and Reference Reagent Program, at a dilution of 1:20,000 in p24 ELISA sample diluent for 1 h at 37°C. Plates were then washed four times with 0.1% Tween 20 in PBS and incubated with goat anti-Human IgG HRP (ThermoFisher, 31412) at a dilution of 1:5000 in p24 ELISA sample diluent. Plates were washed four times with 0.1% Tween 20 in PBS and colorimetric analysis was performed using the TMB Substrate Kit (Thermo Scientific). 30 min later or less reactions were stopped with100 µL of 4 N H_2_SO_4_ and absorbance immediately read at 450 nm using a microplate spectrophotometer. Recombinant p24 was used for the standard curve and sensitive to less than 40 pg of p24.

### Flow Cytometric Detection and Quantitation of HIV-Infected iMG

Intracellular HIV p24 staining in iPSC-derived microglia (iMG) was performed using the BD Cytofix/Cytoperm Fixation/Permeabilization Kit (Cat. No. 554714) following manufacturer recommendations. Briefly, iMGs were isolated and blocked in MACS buffer supplemented with 10 µg/mL human IgG for 30 minutes on ice to reduce nonspecific Fc binding. Cells were then fixed and permeabilized using Cytofix/Cytoperm solution for 20 minutes at 4 °C. Fixed cells were washed twice with Perm/Wash buffer, resuspended in Perm/Wash buffer containing anti-HIV-1 p24 antibody (KC57-FITC, 1:500 dilution), and incubated for 1 hour at 4 °C in the dark. Samples were then washed twice, resuspended in MACS buffer, and immediately analyzed by flow cytometry to quantify p24-positive cells relative to uninfected controls. Data was acquired on a BD FACSCanto II and analyzed using FlowJo 10.10.0 software.

### RNA Isolation from iMG and MDM

Cellular total RNA was isolated from 5 × 10^5^ cells using the RNeasy Mini Kit (Qiagen, 74104). Briefly, cells were pelleted, washed, and lysed in RLT buffer prior to centrifugation through a QIAshredder cell-lysate homogenizer (Qiagen, 79654). Samples were further DNase treated according to manufacturer’s instructions. RNA quality control was performed using an Advanced Analytical Technologies, Inc. (AATI) Fragment Analyzer and integrity RIN/RQN values exceeded 9.5.

### Quantitative Real-time PCR (RT-qPCR)

RNA was isolated from cell pellets using a RNeasy Mini Kit (Qiagen). The RNA was quantified using a NanoDrop Microvolume Spectrophotometer (Thermo Fisher Scientific) and stored at – 80 °C. cDNA synthesis was performed using 1 µg of RNA and the SuperScript IV VILO Master Mix with ezDNase Enzyme (Thermo Fisher Scientific). qPCR amplification reaction was performed using TaqMan probes in a 10 µl reaction using an Applied Biosystems QuantStudio 3 Real-Time PCR System. The specific genes probed, and TaqMan Assay IDs are listed in S1 Table. Reactions were performed in triplicate for all genes. Quantification was performed using the comparative CT method using formula 2^−ΔΔCT^. Relative fold expression was normalized to levels in uninfected iMGs and MDMs for all genes assayed.

### Quantitation of Cytokines in Culture Supernatants by ELISA

iMG and MDM culture supernatant concentrations of IFNγ, TNFα, IL-6, IL-1β and IL-12p70 were determined by enzyme-linked immunosorbent assay (ELISA) using a Luminex assay (R&D Systems) according to manufacturer’s protocol. Briefly, in a 96 well black plate, 50 µL sample in duplicate was incubated with 50 µL antibody coated beads for 2 hours RT on a plate shaker. Plates were then washed 3 times using the BioTek 405 TS (BioTek) and 50 µL of secondary antibody was added and incubated at room temperature for 1 hour on while shaking. Plates were then washed again, 3 times. Finally, 50 µL of streptavidin-RPE was added directly to the secondary antibody and incubated for 30 minutes at room temperature with shaking. Plates were then washed 3 more times and 100 µL of wash buffer was added. Plates were shaken for 5 minutes and then read using Luminex technology on the Milliplex Analyzer (Millipore Sigma). Concentrations were calculated from standard curves using recombinant proteins and expressed in pg/ml.

### Library Preparation, Sequencing and Analysis of Bulk RNA seq

450 ng of RNA per sample was used to construct cDNA libraries via Illumina TruSeq mRNA standard protocols. Each sample was subsequently sequenced on an Illumina NovaSeq 6000 at a depth of approximately 80 million reads per sample. Raw reads were processed using the nf-core/rnaseq pipeline version 3.4 [https://dx.doi.org/10.1038/s41587-020-0439-x]. Briefly, adapter sequences and low-quality reads were filtered and trimmed using FastQ and Trim Galore. Filtered reads were mapped to the human GRCh38 reference genome version 108 using STAR [10.1093/bioinformatics/bts635]. Duplicate reads were removed using MarkDuplicates. Next, Salmon [10.1038/nmeth.4197] was used to generate a count matrix from genes with mapped reads. The full description of the pipeline is available at https://nf-co.re/rnaseq. Gene-level counts were filtered to remove genes with a median expression less than 5. Differential gene expression pre-and post-infection timepoints were evaluated was performed using the DESeq2 R package [https://doi.org/10.1186/s13059-014-0550-8], comparing post-infection timepoint samples to their corresponding cell type (iMG or MDM) uninfected samples. The fgsea R package [10.1101/060012] was used for gene enrichment analysis using the Hallmark gene set collection, using genes ranked by the Wald statistic as reported by DESeq2. Heatmaps, dotplots, and PCA plots were generated using ComplexHeatmap [10.1093/bioinformatics/btw313] and ggplot2 [isbn: 978-3-319-24277-4] R packages. Statistical tests were performed in R. A detailed quantification of the figure data is provided in S4 Table. All datasets generated in this study have been uploaded to the GEO database and are included in S5 Table. Published datasets from the GEO database that we utilized in our analysis are also listed in S5 Table.

### Single cell RNAseq of HIV BaL-infected iMG

Human iPSC-derived microglia (iMG) were generated and matured using standard differentiation protocols. Cells were plated at 0.72 x 105 cells/cm2 on human Fibronectin-coated 12-well plates and cultured at 37 °C/5% CO2 in iMG maintenance media. iMG were infected overnight with HIV BaL at an MOI of 0.25. Uninfected controls underwent identical handling without virus with samples collected at day 0 (baseline/uninfected), day 1, and day 6 post-infection. All work with infectious material was conducted under institutional biosafety approvals and appropriate containment; samples were chemically fixed per 10x guidance prior to removal from containment. Cell fixation was performed following 10x Genomics Cell Fixation Protocol (CG0000776 Rev C). Briefly, cells were pelleted (500 ×g, 5 min), resuspended in Fixation Solution on ice for the recommended duration, then washed and stored at −80 °C. Immediately prior to GEM generation, fixed cells were rehydrated using the protocol-specified Rehydration Buffer, filtered (35 µm), and counted. For each library, we aimed to recover ∼20,000 cells. Libraries were prepared with the Chromium GEM-X Single Cell 3’ v4 Gene Expression chemistry following CG000731. Indexed libraries were pooled equimolarly and sequenced on an Illumina NovaSeq using recommended read configurations for 10x 3′ v4 chemistry, targeting ∼50,000 read pairs per cell. All datasets generated in this study have been uploaded to the GEO database and are included in S5 Table.

### Ethics statement

Human blood used for MDM preparation in this study was obtained from volunteer donors and de-identified before investigator handling. Participants provided informed consent under a protocol approved by the Cincinnati Children’s Hospital Institutional Review Board.

### Code availability

A markdown file and the code used to create all applicable figures are available from the Hagan laboratory repository: httys://haganlab.org/hiv_hand/generic.html. The code is available under an open-source MIT license.

## Supplemental Figures

**S1 FIG.**
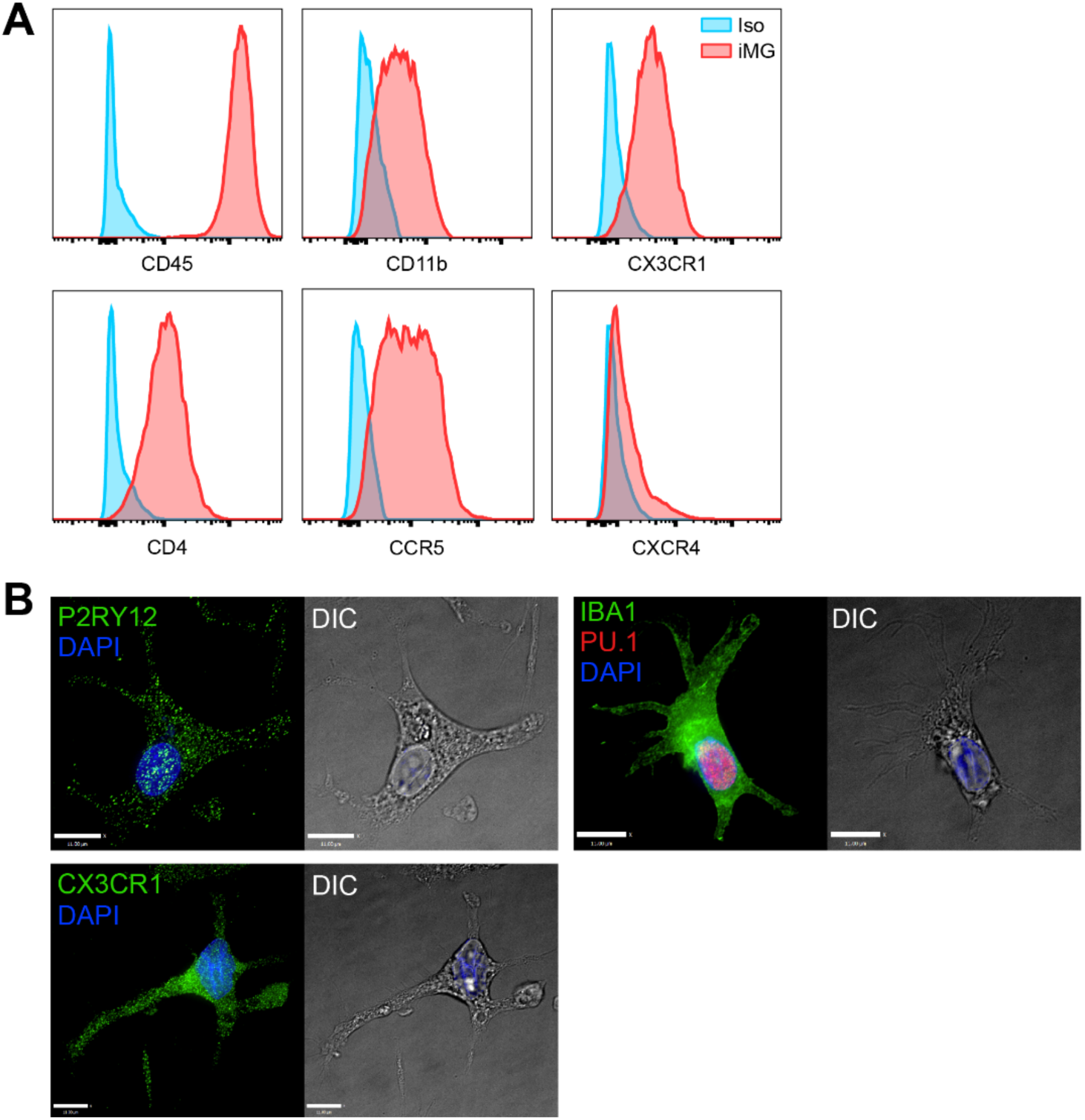
Flow cytometric and immunofluorescence validation of microglial marker expression in iMG. **(A)** Flow panel characterizing surface marker expression (CD45, CD11b, CX3CR1, CD4, CCR5, and CXCR4) in iMG. Blue histograms represent staining with isotype-control antibodies, while red histograms indicate specific staining for the markers listed above. Panels are organized by marker labeling at the bottom of each histogram. (B) Immunofluorescence imaging for microglia markers. Cells were fixed, permeabilized, stained for the microglia markers P2RY12, IBA1, CX3CR1 (green) and the myeloid marker PU.1 (red). The nuclei were counterstained using DAPI (blue). Differential interference contrast (DIC) images confirm characteristic in vitro microglial morphology. Scale bars = 20 µm.

**S2 FIG.**
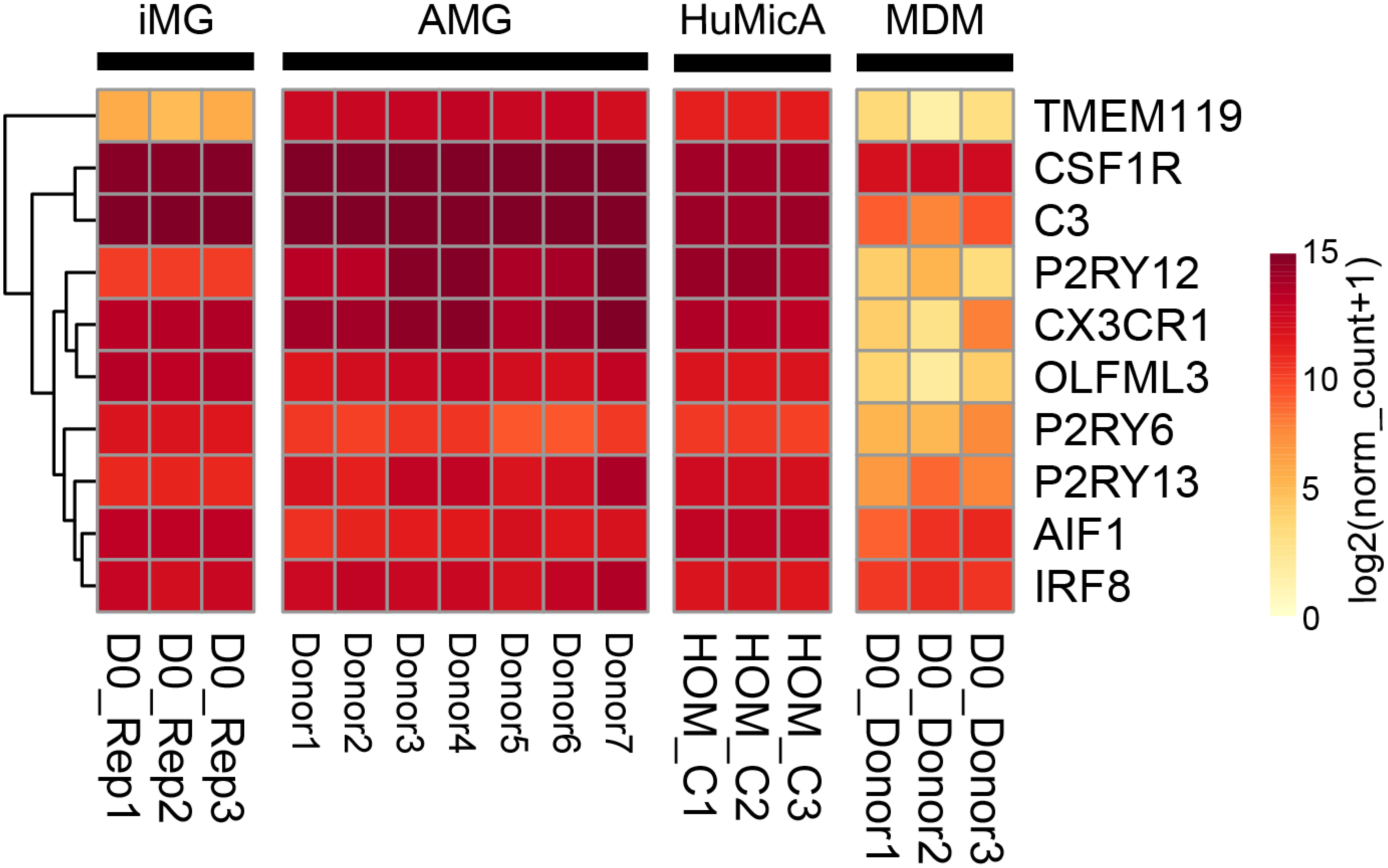
Core Microglial Signature Gene Expression Is Conserved Between iMGs and Adult Human Microglia. Baseline gene expression levels for ten core microglia signature genes in iMG (n=3 biological replicates), adult microglia (AMG), homeostatic adult microglia clusters from the Human Microglia Atlas (HuMicA), and MDMs (n=3 donors). Color in heatmap represents log_2_ (normalized counts+1).

**S3 FIG.**
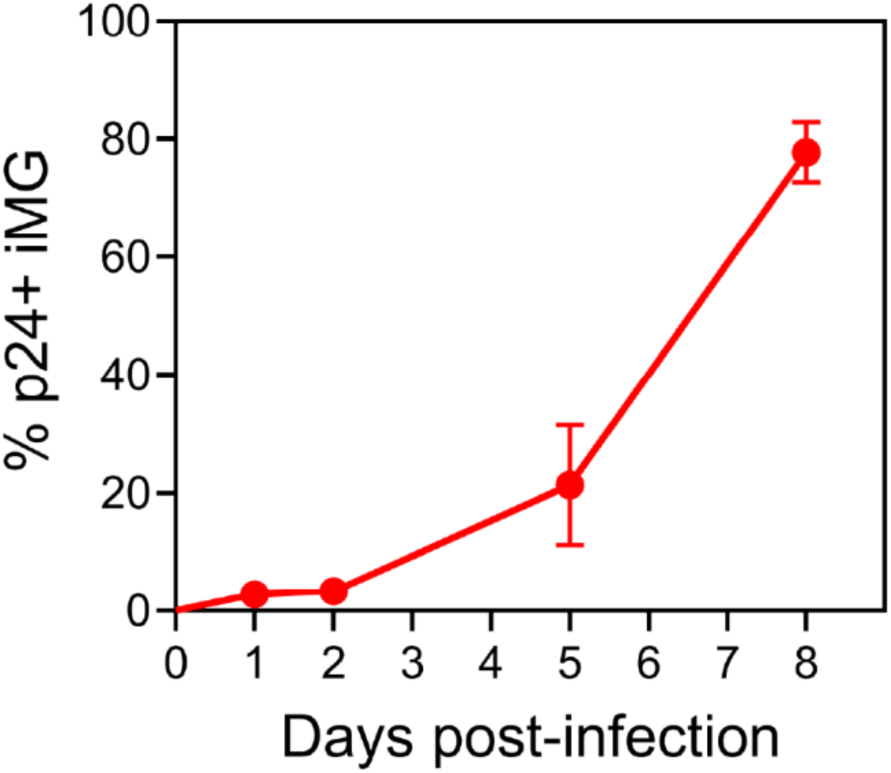
Temporal dynamics of HIV BaL replication in iMG. Percentage of p24⁺ iMG across an 8-day HIV BaL infection time course. Values represent mean ± SD from three from biological replicates. Intracellular p24 was quantified by flow cytometry to determine the proportion of productively infected cells.

**S4 FIG.**
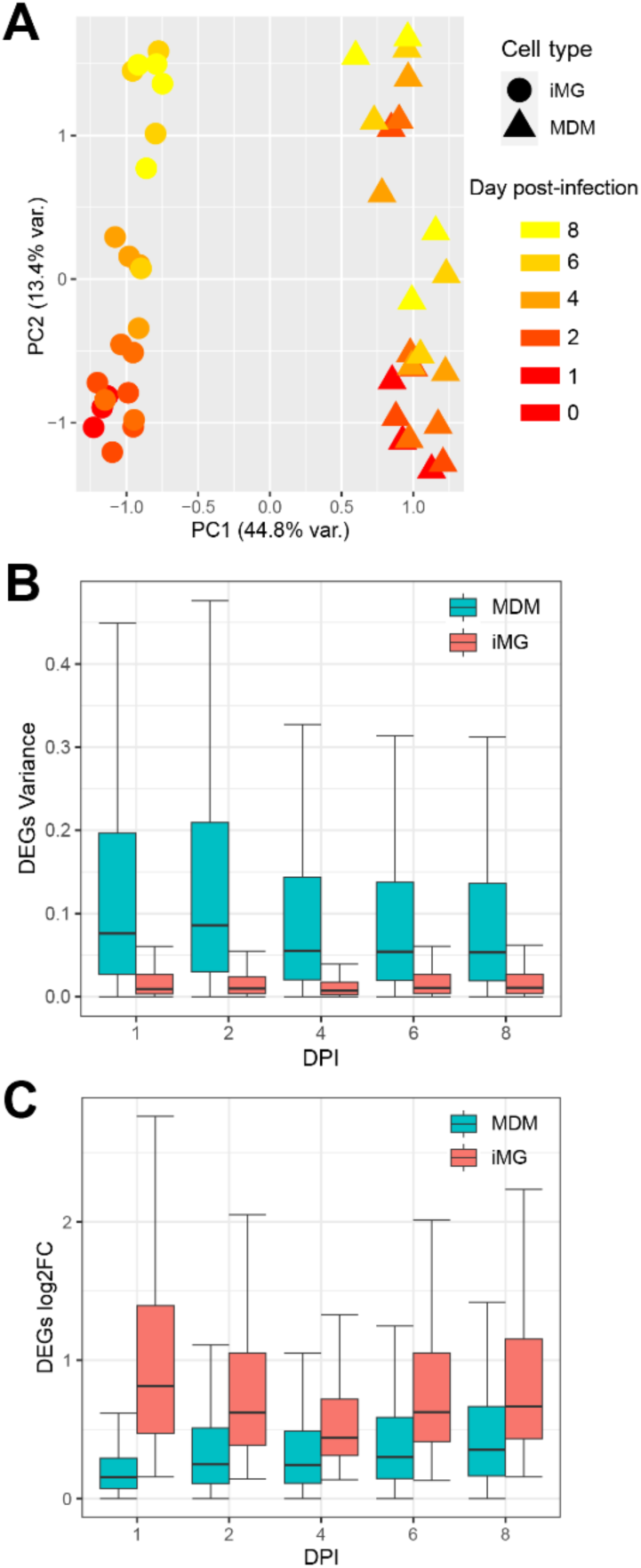
Transcriptomic profiling of HIV-infected microglia and macrophages reveal baseline cell-type divergence and a shared transcriptional trajectory. **(A)** Principal component analysis of global gene expression from HIV-infected iMG (circles) and MDMs (triangles) across an 8-day time course. RNA sequencing was performed on samples from day 0 (uninfected) and days 1, 2, 4, 6 and 8 post-infection. Color indicates days post-infection. **(B)** Quantification of variance in infection induced DEGs from iMGs in both iMG and MDM. **(C)** Quantification of fold changes in infection induced DEGs from iMGs in both iMG and MDM.

**S5 FIG.**
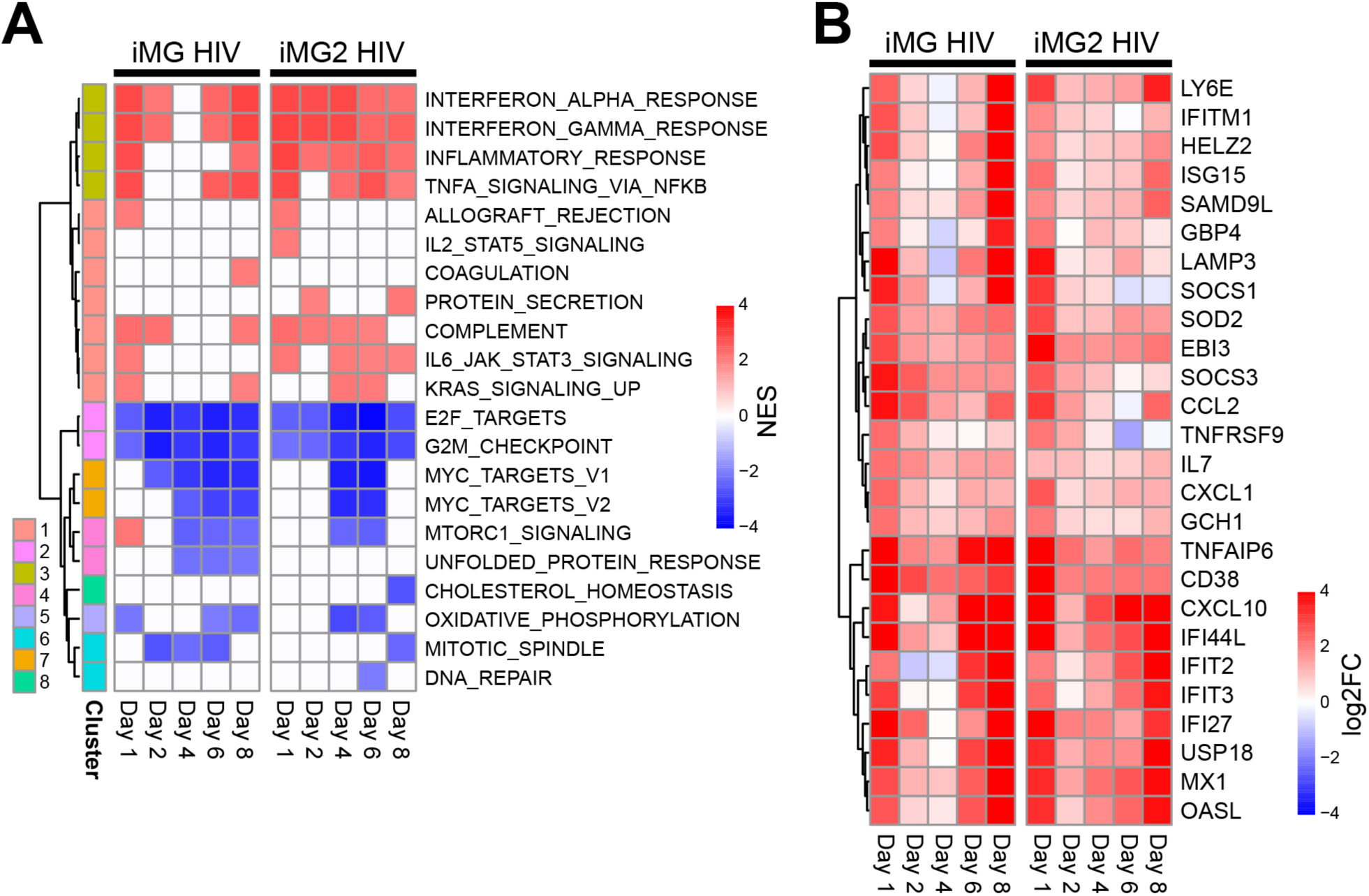
Alternative healthy human iPSC line used to create iMG recreates temporal hyper-inflammatory responses early in HIV-infection. **(A)** Hallmark gene set enrichment analysis of two stem-derived microglia, iMG and iMG2, infected with HIV BaL over an 8-day time course. Color represents normalized enrichment score (NES) in significantly enriched gene sets (FDR<0.05, absolute NES>2). **(B)** Heatmap displaying temporal gene expression subset from Hallmark enrichment analysis pathways taken from cluster 3, Fig 3A (25% overlap, FC≥1).

**S6 FIG.**
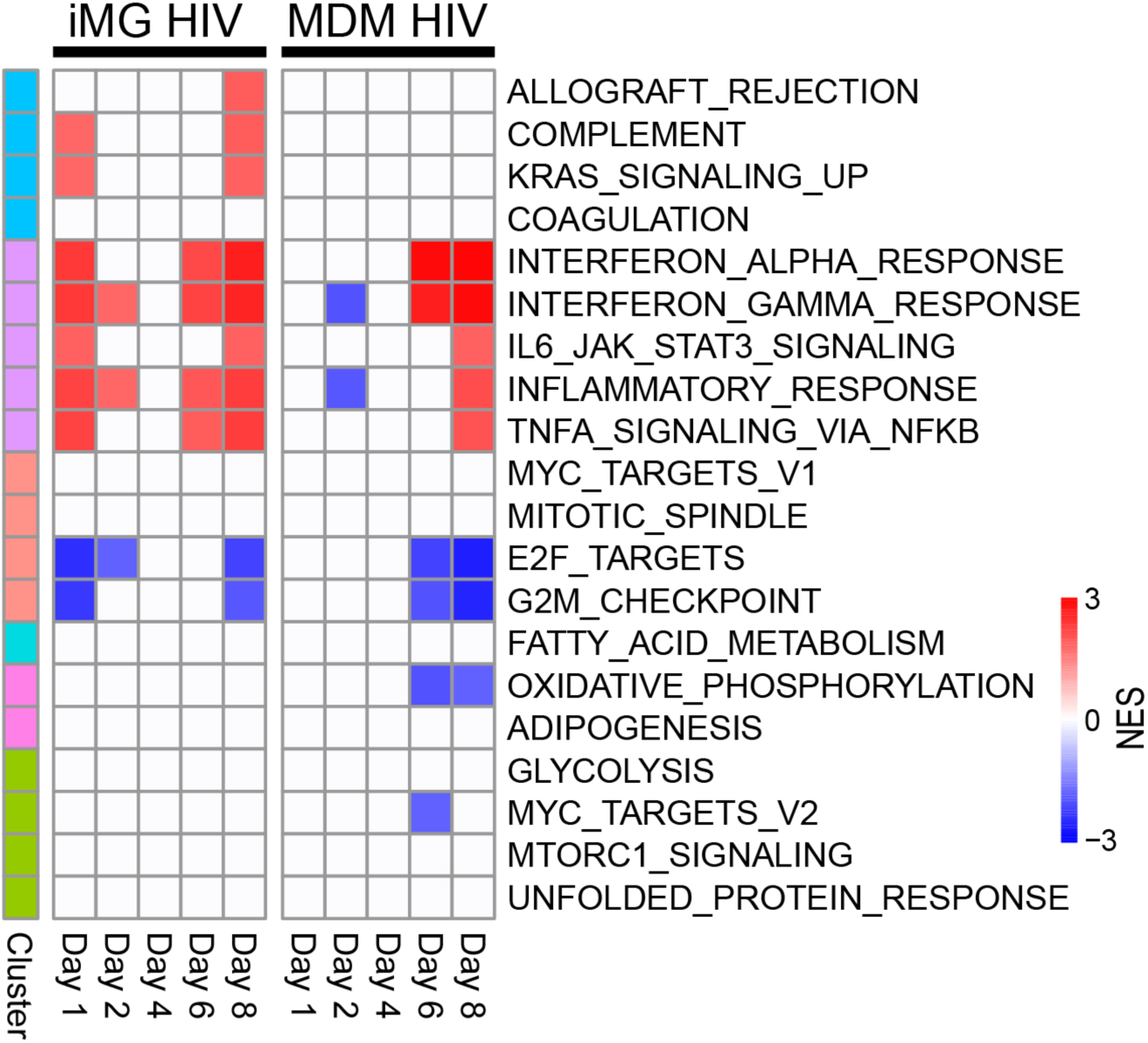
Hyperinflammatory profile of iMG persists after accounting for culture-associated transcriptomic changes in the absence of HIV. Hallmark gene set enrichment analysis of iMG and MDM infected with HIV BaL over an 8-day time course compared to same day uninfected cells. Color represents NES of significantly enriched pathways (FDR<0.05, NES>2).

**S7 FIG.**
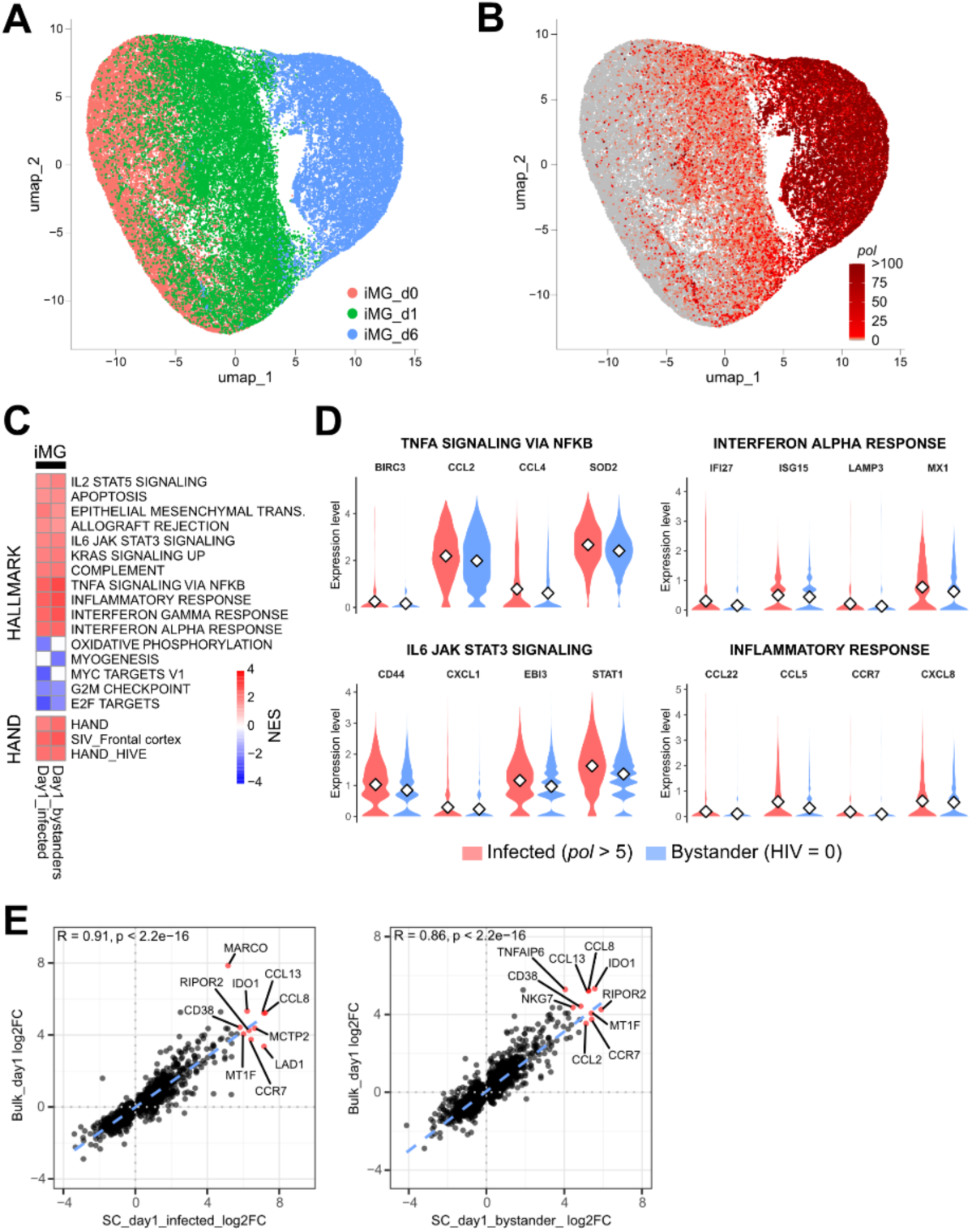
Integrated bulk and single-cell transcriptomic analysis of HIV-1 infection in human microglia. (A, **B)** UMAP projections of scRNA-seq profiles from iPSC-derived microglia (iMG) at day 0 (uninfected), day 1, and day 6 following HIV-1 infection. **(A)** Cells colored by time point. **(B)** Cells colored by HIV *pol* RNA abundance. **(C)** Gene set enrichment analysis of Hallmark and HAND-associated pathways derived from scRNA-seq data for day-1 infected and day-1 bystander iMG populations, shown as normalized enrichment scores (NES). **(D)** Violin plots of SCT-normalized single-cell expression for representative genes from inflammation-related Hallmark pathways in day-1 HIV-infected (red) and bystander (blue) microglia; white diamonds indicate mean expression. Genes shown were selected by ranking bulk RNA-seq DEGs by log₂ fold change and filtering for genes exhibiting consistent induction in scRNA-seq (log₂FC > 0.5). **(E)** Scatter plots comparing bulk RNA-seq and single-cell pseudobulk differential expression (log₂ fold change) for day-1 infected and day-1 bystander infected conditions, with top-ranked genes labeled and Spearman correlation coefficients shown.

**S8 FIG.**
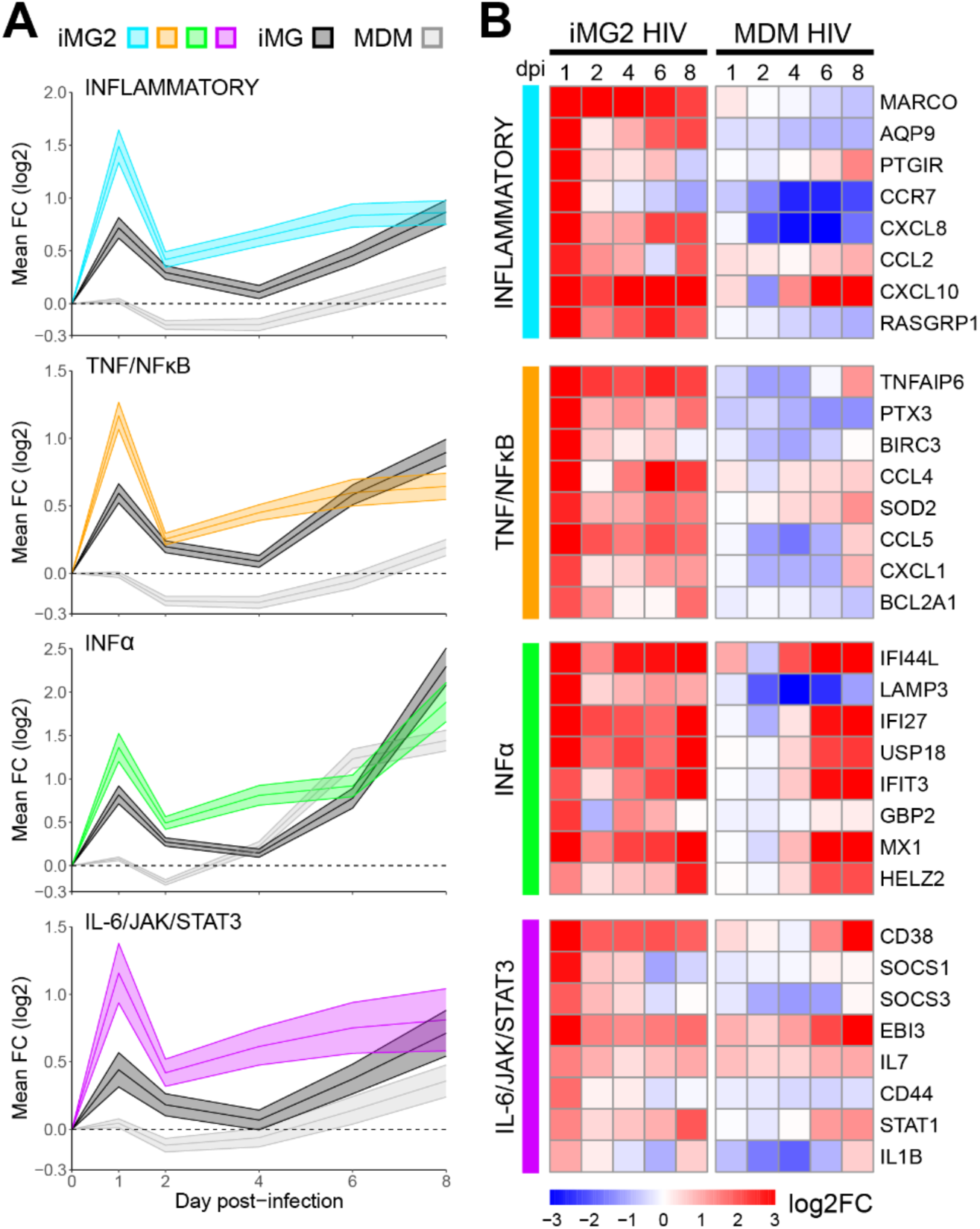
HIV infection induces acute hyper-inflammatory response in iMG2 similar to that observed in iMG1. **(A)** Line graphs of average log_2_FC of all genes present in each inflammation-related pathway across 8-day HIV-infection time course in iMG2. Shaded area represents the 95% confidence interval. Responses in iMG are shown in black and MDMs are shown in gray. **(B)** Heatmap of temporal changes within iMG2 and MDMs in genes among selected inflammation-related Hallmark gene sets significantly enriched post-HIV infection as presented previously (Fig 3A). Top 8 genes with the greatest magnitude of expression changes on day 1 in iMG are represented from each pathway.

**S9 FIG.**
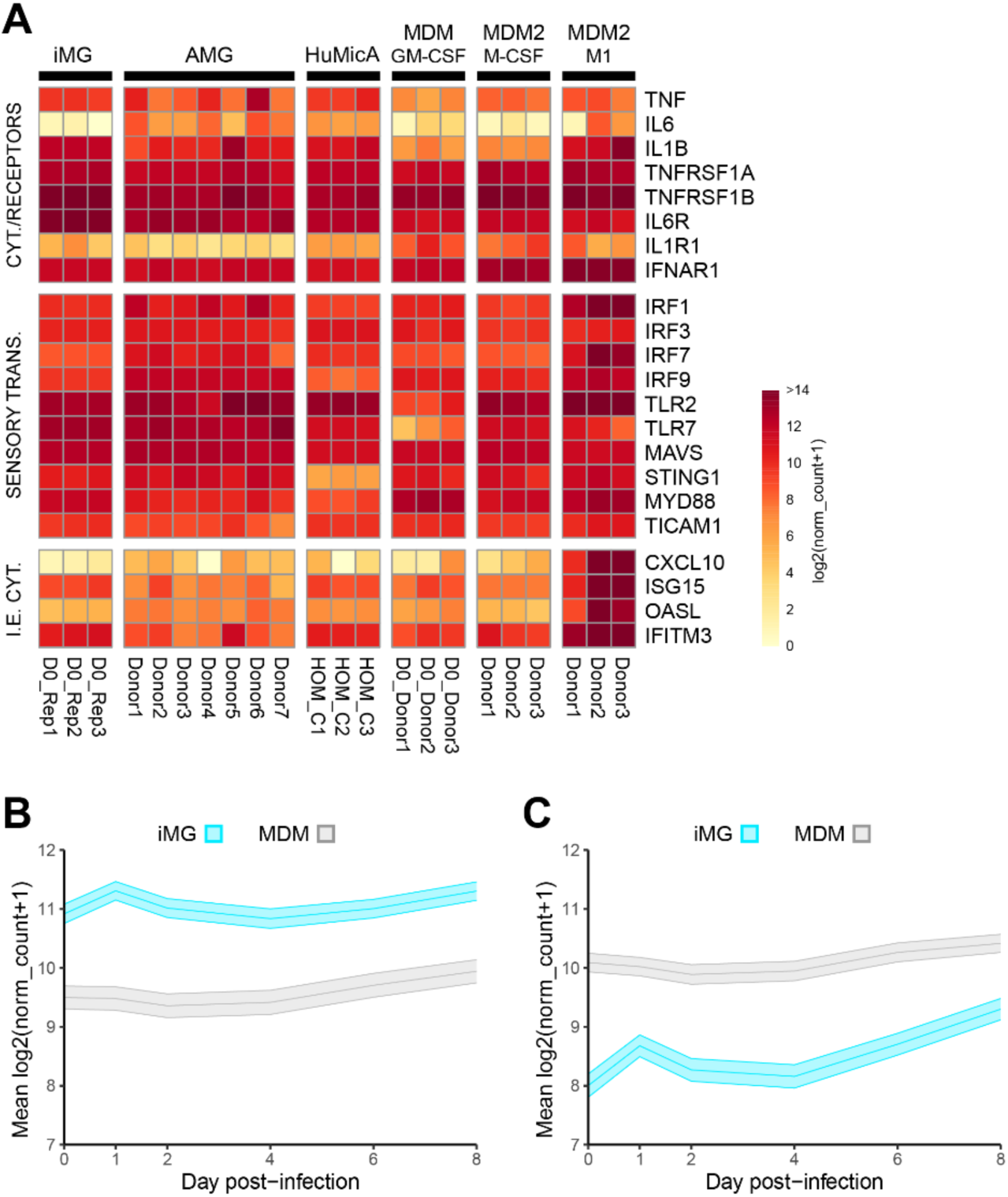
HIV-induced acute hyper-inflammation in iMG is independent of baseline activation state. **(A)** Baseline gene expression for a panel of inflammatory mediators, including inflammatory myeloid cytokines and cognate receptors (CYT./RECEPTORS), pathogen sensors and signal transducers (SENSORY TRANS.) and immediate/early effector cytokines (I.E. CYT). Populations represented are iMG (n=3 biological replicates), adult microglia (AMG), homeostatic adult microglia from the Human Microglia Atlas (HuMicA), human monocyte-derived macrophages matured in GM-CSF (MDM/GM-CSF, n=3 donors), human monocyte-derived macrophages matured in M-CSF (MDM2/M-CSF, n=3 donors) and human monocyte-derived macrophages stimulated with LPS to induce an M1 phenotype (MDM2/M1, n=3 donors). Color in heatmap represents log_2_ (normalized counts+1). **(B)** Line graphs of mean log2FC of genes expressed higher in iMG at baseline vs. MDM from four merged inflammation-related Hallmark pathways (Inflammatory, TNF/NFκB, IFNα, IL-6/JAK/STAT3, n=421 genes) across 8-day HIV-infection time course. Shaded area represents the 95% confidence interval. **(C)** Line graphs of mean log2FC of genes expressed higher in MDM at baseline vs. iMG from four merged inflammation-related Hallmark pathways (Inflammatory, TNF/NFκB, IFNα, IL-6/JAK/STAT3, n=421 genes) across 8-day HIV-infection time course. Shaded area represents the 95% confidence interval.

**S10 FIG.**
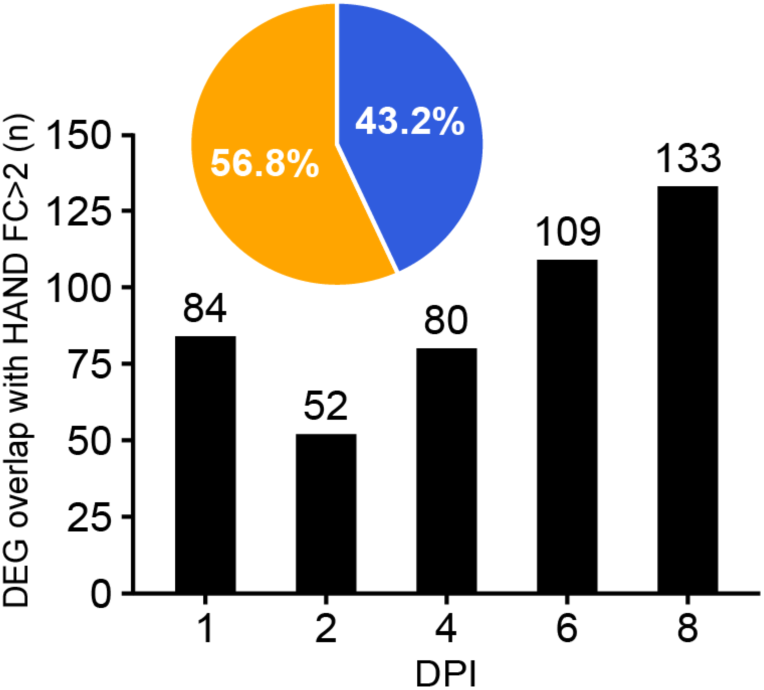
HIV-infected iMGs show gene expression changes linked to HAND. Graph illustrating DEGs from HIV-infected iMGs on specific days post-infection that are part of the Borjabad HAND signature. Pie chart represents intersection of DEGs from infected iMGs with the Borjabad HAND signature, where overlap is represented in orange, absence in blue, percentages listed in white.

## Supporting Information Captions

### Table captions

**S1 Table. Reagents.** Excel worksheet containing details of reagents, media formulations, and TaqMan qPCR probes utilized in this manuscript.

**S2 Table. RNAseq counts.** Excel worksheet containing raw and normalized counts for samples analyzed in this manuscript.

**S3 Table. HIV Infection Time Course Differentially Expressed Genes (DEGs).** Lists of differentially expressed genes (DEGs) for HIV-infected iMG and MDM across the infection time course.

**S4 Table. Figure Quantification.** Excel worksheet containing figure quantification data, with individual sheets corresponding to the quantification results for each specific figure panel.

**S5 Table. GEO Accession Numbers.** File contains GEO accession numbers corresponding to the samples used for RNAseq analysis in this manuscript.

## References

1. Organization WH. The Global Health Observatory 2024 [cited 2026 July 2]. Available from: https://www.who.int/data/gho/data/themes/hiv-aids.

2. McArthur JC, Steiner J, Sacktor N, Nath A. Human immunodeficiency virus-associated neurocognitive disorders: Mind the gap. Ann Neurol. 2010;67(6):699–714. doi: 10.1002/ana.22053. PubMed PMID: 20517932.

3. Ojeda-Juarez D, Kaul M. Transcriptomic and Genetic Profiling of HIV-Associated Neurocognitive Disorders. Front Mol Biosci. 2021;8:721954. Epub 20211029. doi: 10.3389/fmolb.2021.721954. PubMed PMID: 34778371; PubMed Central PMCID:PMC8586712.

4. Saylor D, Dickens AM, Sacktor N, Haughey N, Slusher B, Pletnikov M, et al. HIV-associated neurocognitive disorder - pathogenesis and prospects for treatment. Nat Rev Neurol. 2016;12(5):309. Epub 20160415. doi: 10.1038/nrneurol.2016.53. PubMed PMID: 27080521; PubMed Central PMCID:PMC5842923.

5. Vella S, Schwartlander B, Sow SP, Eholie SP, Murphy RL. The history of antiretroviral therapy and of its implementation in resource-limited areas of the world. AIDS. 2012;26(10):1231–41. doi: 10.1097/QAD.0b013e32835521a3. PubMed PMID: 22706009.

6. Tang Y, Chaillon A, Gianella S, Wong LM, Li D, Simermeyer TL, et al. Brain microglia serve as a persistent HIV reservoir despite durable antiretroviral therapy. Journal of Clinical Investigation. 2023;133(12). doi: 10.1172/jci167417.

7. Norris GT, Kipnis J. Immune cells and CNS physiology: Microglia and beyond. J Exp Med. 2019;216(1):60-70. Epub 20181130. doi: 10.1084/jem.20180199. PubMed PMID: 30504438; PubMed Central PMCID:PMC6314530.

8. Abreu CM, Veenhuis RT, Avalos CR, Graham S, Parrilla DR, Ferreira EA, et al. Myeloid and CD4 T Cells Comprise the Latent Reservoir in Antiretroviral Therapy-Suppressed SIVmac251-Infected Macaques. mBio. 2019;10(4). Epub 20190820. doi: 10.1128/mBio.01659-19. PubMed PMID: 31431552; PubMed Central PMCID:PMC6703426.

9. Crowe S, Zhu T, Muller WA. The contribution of monocyte infection and trafficking to viral persistence, and maintenance of the viral reservoir in HIV infection. J Leukoc Biol. 2003;74(5):635–41. Epub 20030821. doi: 10.1189/jlb.0503204. PubMed PMID: 12960232.

10. Honeycutt JB, Liao B, Nixon CC, Cleary RA, Thayer WO, Birath SL, et al. T cells establish and maintain CNS viral infection in HIV-infected humanized mice. J Clin Invest. 2018;128(7):2862–76. Epub 20180604. doi: 10.1172/JCI98968. PubMed PMID: 29863499; PubMed Central PMCID:PMC6026008.

11. Joseph SB, Arrildt KT, Sturdevant CB, Swanstrom R. HIV-1 target cells in the CNS. J Neurovirol. 2015;21(3):276–89. Epub 20140919. doi: 10.1007/s13365-014-0287-x. PubMed PMID: 25236812; PubMed Central PMCID:PMC4366351.

12. Mitchell BI, Laws EI, Ndhlovu LC. Impact of Myeloid Reservoirs in HIV Cure Trials. Curr HIV/AIDS Rep. 2019;16(2):129–40. doi: 10.1007/s11904-019-00438-5. PubMed PMID: 30835045; PubMed Central PMCID:PMC6589351.

13. Trillo-Pazos G, Diamanturos A, Rislove L, Menza T, Chao W, Belem P, et al. Detection of HIV-1 DNA in microglia/macrophages, astrocytes and neurons isolated from brain tissue with HIV-1 encephalitis by laser capture microdissection. Brain Pathol. 2003;13(2):144–54. doi: 10.1111/j.1750-3639.2003.tb00014.x. PubMed PMID: 12744468; PubMed Central PMCID:PMC8096041.

14. Tang Y, Jiang G. Eradication of human immunodeficiency virus-1 reservoir in the brain microglia. Neural Regen Res. 2023;18(3):552–3. doi: 10.4103/1673-5374.350198. PubMed PMID: 36018174; PubMed Central PMCID:PMC9727433.

15. Spudich S, Gonzalez-Scarano F. HIV-1-related central nervous system disease: current issues in pathogenesis, diagnosis, and treatment. Cold Spring Harb Perspect Med. 2012;2(6):a007120. doi: 10.1101/cshperspect.a007120. PubMed PMID: 22675662; PubMed Central PMCID:PMC3367536.

16. Loewenstein RJ, Sharfstein SS. Neuropsychiatric aspects of acquired immune deficiency syndrome. Int J Psychiatry Med. 1983;13(4):255–60. doi: 10.2190/repy-d2wd-07nm-mbrt. PubMed PMID: 6671858.

17. Navia BA, Cho ES, Petito CK, Price RW. The AIDS dementia complex: II. Neuropathology. Ann Neurol. 1986;19(6):525–35. doi: 10.1002/ana.410190603. PubMed PMID: 3014994.

18. Navia BA, Jordan BD, Price RW. The AIDS dementia complex: I. Clinical features. Ann Neurol. 1986;19(6):517–24. doi: 10.1002/ana.410190602. PubMed PMID: 3729308.

19. Borjabad A, Volsky DJ. Common transcriptional signatures in brain tissue from patients with HIV-associated neurocognitive disorders, Alzheimer’s disease, and Multiple Sclerosis. J Neuroimmune Pharmacol. 2012;7(4):914–26. Epub 2012/10/16. doi: 10.1007/s11481-012-9409-5. PubMed PMID: 23065460; PubMed Central PMCID:PMC3515772.

20. Ellis R, Langford D, Masliah E. HIV and antiretroviral therapy in the brain: neuronal injury and repair. Nat Rev Neurosci. 2007;8(1):33–44. doi: 10.1038/nrn2040. PubMed PMID: 17180161.

21. Sacktor N, McDermott MP, Marder K, Schifitto G, Selnes OA, McArthur JC, et al. HIV-associated cognitive impairment before and after the advent of combination therapy. J Neurovirol. 2002;8(2):136–42. doi: 10.1080/13550280290049615. PubMed PMID: 11935465.

22. Harezlak J, Buchthal S, Taylor M, Schifitto G, Zhong J, Daar E, et al. Persistence of HIV-associated cognitive impairment, inflammation, and neuronal injury in era of highly active antiretroviral treatment. AIDS. 2011;25(5):625–33. doi: 10.1097/QAD.0b013e3283427da7. PubMed PMID: 21297425; PubMed Central PMCID:PMC4326227.

23. Heaton RK, Clifford DB, Franklin DR, Jr., Woods SP, Ake C, Vaida F, et al. HIV-associated neurocognitive disorders persist in the era of potent antiretroviral therapy: CHARTER Study. Neurology. 2010;75(23):2087–96. doi: 10.1212/WNL.0b013e318200d727. PubMed PMID: 21135382; PubMed Central PMCID:PMC2995535.

24. Simioni S, Cavassini M, Annoni JM, Rimbault Abraham A, Bourquin I, Schiffer V, et al. Cognitive dysfunction in HIV patients despite long-standing suppression of viremia. AIDS. 2010;24(9):1243–50. doi: 10.1097/QAD.0b013e3283354a7b. PubMed PMID: 19996937.

25. Kwon HS, Koh SH. Neuroinflammation in neurodegenerative disorders: the roles of microglia and astrocytes. Transl Neurodegener. 2020;9(1):42. Epub 20201126. doi: 10.1186/s40035-020-00221-2. PubMed PMID: 33239064; PubMed Central PMCID:PMC7689983.

26. Muzio L, Viotti A, Martino G. Microglia in Neuroinflammation and Neurodegeneration: From Understanding to Therapy. Front Neurosci. 2021;15:742065. Epub 20210924. doi: 10.3389/fnins.2021.742065. PubMed PMID: 34630027; PubMed Central PMCID:PMC8497816.

27. Abud EM, Ramirez RN, Martinez ES, Healy LM, Nguyen CHH, Newman SA, et al. iPSC-Derived Human Microglia-like Cells to Study Neurological Diseases. Neuron. 2017;94(2):278–93 e9. Epub 2017/04/21. doi: 10.1016/j.neuron.2017.03.042. PubMed PMID: 28426964; PubMed Central PMCID:PMC5482419.

28. McQuade A, Coburn M, Tu CH, Hasselmann J, Davtyan H, Blurton-Jones M. Development and validation of a simplified method to generate human microglia from pluripotent stem cells. Mol Neurodegener. 2018;13(1):67. Epub 2018/12/24. doi: 10.1186/s13024-018-0297-x. PubMed PMID: 30577865; PubMed Central PMCID:PMC6303871.

29. Rai MA, Hammonds J, Pujato M, Mayhew C, Roskin K, Spearman P. Comparative analysis of human microglial models for studies of HIV replication and pathogenesis. Retrovirology. 2020;17(1):35. Epub 20201119. doi: 10.1186/s12977-020-00544-y. PubMed PMID: 33213476; PubMed Central PMCID:PMC7678224.

30. Galatro TF, Holtman IR, Lerario AM, Vainchtein ID, Brouwer N, Sola PR, et al. Transcriptomic analysis of purified human cortical microglia reveals age-associated changes. Nat Neurosci. 2017;20(8):1162–71. Epub 20170703. doi: 10.1038/nn.4597. PubMed PMID: 28671693.

31. Gosselin D, Skola D, Coufal NG, Holtman IR, Schlachetzki JCM, Sajti E, et al. An environment-dependent transcriptional network specifies human microglia identity. Science. 2017;356(6344). Epub 2017/05/27. doi: 10.1126/science.aal3222. PubMed PMID: 28546318; PubMed Central PMCID:PMC5858585.

32. Muffat J, Li Y, Yuan B, Mitalipova M, Omer A, Corcoran S, et al. Efficient derivation of microglia-like cells from human pluripotent stem cells. Nat Med. 2016;22(11):1358–67. Epub 20160926. doi: 10.1038/nm.4189. PubMed PMID: 27668937; PubMed Central PMCID:PMC5101156.

33. Martins-Ferreira R, Calafell-Segura J, Leal B, Rodriguez-Ubreva J, Martinez-Saez E, Mereu E, et al. The Human Microglia Atlas (HuMicA) unravels changes in disease-associated microglia subsets across neurodegenerative conditions. Nat Commun. 2025;16(1):739. Epub 20250116. doi: 10.1038/s41467-025-56124-1. PubMed PMID: 39820004; PubMed Central PMCID:PMC11739505.

34. Oldham MC, Konopka G, Iwamoto K, Langfelder P, Kato T, Horvath S, et al. Functional organization of the transcriptome in human brain. Nat Neurosci. 2008;11(11):1271–82. Epub 20081012. doi: 10.1038/nn.2207. PubMed PMID: 18849986; PubMed Central PMCID:PMC2756411.

35. Patir A, Shih B, McColl BW, Freeman TC. A core transcriptional signature of human microglia: Derivation and utility in describing region-dependent alterations associated with Alzheimer’s disease. Glia. 2019;67(7):1240–53. Epub 2019/02/14. doi: 10.1002/glia.23572. PubMed PMID: 30758077.

36. Wehrspaun CC, Haerty W, Ponting CP. Microglia recapitulate a hematopoietic master regulator network in the aging human frontal cortex. Neurobiol Aging. 2015;36(8):2443 e9- e20. Epub 20150425. doi: 10.1016/j.neurobiolaging.2015.04.008. PubMed PMID: 26002684; PubMed Central PMCID:PMC4503803.

37. Zhang Y, Sloan SA, Clarke LE, Caneda C, Plaza CA, Blumenthal PD, et al. Purification and Characterization of Progenitor and Mature Human Astrocytes Reveals Transcriptional and Functional Differences with Mouse. Neuron. 2016;89(1):37–53. Epub 20151210. doi: 10.1016/j.neuron.2015.11.013. PubMed PMID: 26687838; PubMed Central PMCID:PMC4707064.

38. Haage V, Semtner M, Vidal RO, Hernandez DP, Pong WW, Chen Z, et al. Comprehensive gene expression meta-analysis identifies signature genes that distinguish microglia from peripheral monocytes/macrophages in health and glioma. Acta Neuropathol Commun. 2019;7(1):20. Epub 20190214. doi: 10.1186/s40478-019-0665-y. PubMed PMID: 30764877; PubMed Central PMCID:PMC6376799.

39. Olah M, Patrick E, Villani AC, Xu J, White CC, Ryan KJ, et al. A transcriptomic atlas of aged human microglia. Nat Commun. 2018;9(1):539. Epub 20180207. doi: 10.1038/s41467-018-02926-5. PubMed PMID: 29416036; PubMed Central PMCID:PMC5803269.

40. Liberzon A, Birger C, Thorvaldsdottir H, Ghandi M, Mesirov JP, Tamayo P. The Molecular Signatures Database (MSigDB) hallmark gene set collection. Cell Syst. 2015;1(6):417–25. doi: 10.1016/j.cels.2015.12.004. PubMed PMID: 26771021; PubMed Central PMCID:PMC4707969.

41. Subramanian A, Tamayo P, Mootha VK, Mukherjee S, Ebert BL, Gillette MA, et al. Gene set enrichment analysis: a knowledge-based approach for interpreting genome-wide expression profiles. Proc Natl Acad Sci U S A. 2005;102(43):15545–50. doi: 10.1073/pnas.0506580102. PubMed PMID: 16199517; PubMed Central PMCID:PMC1239896.

42. Li S, Rouphael N, Duraisingham S, Romero-Steiner S, Presnell S, Davis C, et al. Molecular signatures of antibody responses derived from a systems biology study of five human vaccines. Nat Immunol. 2014;15(2):195–204. Epub 20131215. doi: 10.1038/ni.2789. PubMed PMID: 24336226; PubMed Central PMCID:PMC3946932.

43. Nakaya HI, Hagan T, Duraisingham SS, Lee EK, Kwissa M, Rouphael N, et al. Systems Analysis of Immunity to Influenza Vaccination across Multiple Years and in Diverse Populations Reveals Shared Molecular Signatures. Immunity. 2015;43(6):1186–98. doi: 10.1016/j.immuni.2015.11.012. PubMed PMID: 26682988; PubMed Central PMCID:PMC4859820.

44. Kong W, Frouard J, Xie G, Corley MJ, Helmy E, Zhang G, et al. Neuroinflammation generated by HIV-infected microglia promotes dysfunction and death of neurons in human brain organoids. PNAS Nexus. 2024;3(5):pgae179. Epub 20240429. doi: 10.1093/pnasnexus/pgae179. PubMed PMID: 38737767; PubMed Central PMCID:PMC11086946.

45. Li H, McLaurin KA, Illenberger JM, Mactutus CF, Booze RM. Microglial HIV-1 Expression: Role in HIV-1 Associated Neurocognitive Disorders. Viruses. 2021;13(5). Epub 20210517. doi: 10.3390/v13050924. PubMed PMID: 34067600; PubMed Central PMCID:PMC8155894.

46. Williams ME, Naude PJW. The relationship between HIV-1 neuroinflammation, neurocognitive impairment and encephalitis pathology: A systematic review of studies investigating post-mortem brain tissue. Rev Med Virol. 2024;34(1):e2519. doi: 10.1002/rmv.2519. PubMed PMID: 38282400; PubMed Central PMCID:PMC10909494.

47. Polyak MJ, Vivithanaporn P, Maingat FG, Walsh JG, Branton W, Cohen EA, et al. Differential type 1 interferon-regulated gene expression in the brain during AIDS: interactions with viral diversity and neurovirulence. FASEB J. 2013;27(7):2829–44. Epub 20130422. doi: 10.1096/fj.13-227868. PubMed PMID: 23608145; PubMed Central PMCID:PMC3955194.

48. Sanfilippo C, Pinzone MR, Cambria D, Longo A, Palumbo M, Di Marco R, et al. OAS Gene Family Expression Is Associated with HIV-Related Neurocognitive Disorders. Mol Neurobiol. 2018;55(3):1905–14. Epub 20170224. doi: 10.1007/s12035-017-0460-3. PubMed PMID: 28236279.

49. Wang S, Ding X, Li Z, Rao F, Xu H, Lu J, et al. Comprehensive analyses identify potential biomarkers for encephalitis in HIV infection. Sci Rep. 2023;13(1):18418. Epub 20231027. doi: 10.1038/s41598-023-45922-6. PubMed PMID: 37891420; PubMed Central PMCID:PMC10611703.

50. Zaritsky LA, Dery A, Leong WY, Gama L, Clements JE. Tissue-specific interferon alpha subtype response to SIV infection in brain, spleen, and lung. J Interferon Cytokine Res. 2013;33(1):24–33. Epub 20121010. doi: 10.1089/jir.2012.0018. PubMed PMID: 23050948; PubMed Central PMCID:PMC3539257.

51. Williams ME, Stein DJ, Joska JA, Naude PJW. Cerebrospinal fluid immune markers and HIV-associated neurocognitive impairments: A systematic review. J Neuroimmunol. 2021;358:577649. Epub 20210630. doi: 10.1016/j.jneuroim.2021.577649. PubMed PMID: 34280844.

52. Guerreiro S, Privat AL, Bressac L, Toulorge D. CD38 in Neurodegeneration and Neuroinflammation. Cells. 2020;9(2). Epub 20200218. doi: 10.3390/cells9020471. PubMed PMID: 32085567; PubMed Central PMCID:PMC7072759.

53. Hammonds JE, Beeman N, Ding L, Takushi S, Francis AC, Wang JJ, et al. Siglec-1 initiates formation of the virus-containing compartment and enhances macrophage-to-T cell transmission of HIV-1. PLoS Pathog. 2017;13(1):e1006181. Epub 20170127. doi: 10.1371/journal.ppat.1006181. PubMed PMID: 28129379; PubMed Central PMCID:PMC5298340.

54. Izquierdo-Useros N, Lorizate M, Puertas MC, Rodriguez-Plata MT, Zangger N, Erikson E, et al. Siglec-1 is a novel dendritic cell receptor that mediates HIV-1 trans-infection through recognition of viral membrane gangliosides. PLoS Biol. 2012;10(12):e1001448. doi: 10.1371/journal.pbio.1001448. PubMed PMID: 23271952; PubMed Central PMCID:PMC3525531 application based on this work has been filed (EP11382392.6, 2011). The authors declare that no other competing financial interests exist.

55. Puryear WB, Akiyama H, Geer SD, Ramirez NP, Yu X, Reinhard BM, et al. Interferon-inducible mechanism of dendritic cell-mediated HIV-1 dissemination is dependent on Siglec-1/CD169. PLoS Pathog. 2013;9(4):e1003291. doi: 10.1371/journal.ppat.1003291. PubMed PMID: 23593001; PubMed Central PMCID:PMC3623718.

56. Malim MH, Bieniasz PD. HIV Restriction Factors and Mechanisms of Evasion. Cold Spring Harb Perspect Med. 2012;2(5):a006940. doi: 10.1101/cshperspect.a006940. PubMed PMID: 22553496; PubMed Central PMCID:PMC3331687.

57. V DU, De Crignis E, Re MC. Host Restriction Factors and Human Immunodeficiency Virus (HIV-1): A Dynamic Interplay Involving All Phases of the Viral Life Cycle. Curr HIV Res. 2018;16(3):184–207. doi: 10.2174/1570162X16666180817115830. PubMed PMID: 30117396.

58. Braun E, Hotter D, Koepke L, Zech F, Gross R, Sparrer KMJ, et al. Guanylate-Binding Proteins 2 and 5 Exert Broad Antiviral Activity by Inhibiting Furin-Mediated Processing of Viral Envelope Proteins. Cell Rep. 2019;27(7):2092–104 e10. doi: 10.1016/j.celrep.2019.04.063. PubMed PMID: 31091448.

59. Laguette N, Sobhian B, Casartelli N, Ringeard M, Chable-Bessia C, Segeral E, et al. SAMHD1 is the dendritic- and myeloid-cell-specific HIV-1 restriction factor counteracted by Vpx. Nature. 2011;474(7353):654–7. Epub 20110525. doi: 10.1038/nature10117. PubMed PMID: 21613998; PubMed Central PMCID:PMC3595993.

60. St Gelais C, Wu L. SAMHD1: a new insight into HIV-1 restriction in myeloid cells. Retrovirology. 2011;8:55. Epub 20110708. doi: 10.1186/1742-4690-8-55. PubMed PMID: 21740548; PubMed Central PMCID:PMC3142215.

61. Neil SJ, Zang T, Bieniasz PD. Tetherin inhibits retrovirus release and is antagonized by HIV-1 Vpu. Nature. 2008;451(7177):425–30. Epub 20080116. doi: 10.1038/nature06553. PubMed PMID: 18200009.

62. Borjabad A, Morgello S, Chao W, Kim SY, Brooks AI, Murray J, et al. Significant effects of antiretroviral therapy on global gene expression in brain tissues of patients with HIV-1-associated neurocognitive disorders. PLoS Pathog. 2011;7(9):e1002213. Epub 2011/09/13. doi: 10.1371/journal.ppat.1002213. PubMed PMID: 21909266; PubMed Central PMCID:PMC3164642.

63. Gelman BB, Chen T, Lisinicchia JG, Soukup VM, Carmical JR, Starkey JM, et al. The National NeuroAIDS Tissue Consortium brain gene array: two types of HIV-associated neurocognitive impairment. PLoS One. 2012;7(9):e46178. Epub 20120926. doi: 10.1371/journal.pone.0046178. PubMed PMID: 23049970; PubMed Central PMCID:PMC3458860.

64. Mavian C, Ramirez-Mata AS, Dollar JJ, Nolan DJ, Cash M, White K, et al. Brain tissue transcriptomic analysis of SIV-infected macaques identifies several altered metabolic pathways linked to neuropathogenesis and poly (ADP-ribose) polymerases (PARPs) as potential therapeutic targets. J Neurovirol. 2021;27(1):101–15. Epub 20210106. doi: 10.1007/s13365-020-00927-z. PubMed PMID: 33405206; PubMed Central PMCID:PMC7786889.

65. Jao J, Ciernia AV. MGEnrichment: A web application for microglia gene list enrichment analysis. PLoS Comput Biol. 2021;17(11):e1009160. Epub 20211117. doi: 10.1371/journal.pcbi.1009160. PubMed PMID: 34788279; PubMed Central PMCID:PMC8598070.

66. Hammond TR, Dufort C, Dissing-Olesen L, Giera S, Young A, Wysoker A, et al. Single-Cell RNA Sequencing of Microglia throughout the Mouse Lifespan and in the Injured Brain Reveals Complex Cell-State Changes. Immunity. 2019;50(1):253–71 e6. Epub 20181121. doi: 10.1016/j.immuni.2018.11.004. PubMed PMID: 30471926; PubMed Central PMCID:PMC6655561.

67. Zhou Y, Song WM, Andhey PS, Swain A, Levy T, Miller KR, et al. Human and mouse single-nucleus transcriptomics reveal TREM2-dependent and TREM2-independent cellular responses in Alzheimer’s disease. Nat Med. 2020;26(1):131–42. Epub 20200113. doi: 10.1038/s41591-019-0695-9. PubMed PMID: 31932797; PubMed Central PMCID:PMC6980793.

68. Duverger A, Wolschendorf F, Anderson JC, Wagner F, Bosque A, Shishido T, et al. Kinase control of latent HIV-1 infection: PIM-1 kinase as a major contributor to HIV-1 reactivation. J Virol. 2014;88(1):364–76. Epub 20131023. doi: 10.1128/JVI.02682-13. PubMed PMID: 24155393; PubMed Central PMCID:PMC3911731.

69. Vicenzi E, Poli G. The interferon-stimulated gene TRIM22: A double-edged sword in HIV-1 infection. Cytokine Growth Factor Rev. 2018;40:40–7. Epub 20180210. doi: 10.1016/j.cytogfr.2018.02.001. PubMed PMID: 29650252.

70. Woods MW, Kelly JN, Hattlmann CJ, Tong JG, Xu LS, Coleman MD, et al. Human HERC5 restricts an early stage of HIV-1 assembly by a mechanism correlating with the ISGylation of Gag. Retrovirology. 2011;8:95. Epub 20111117. doi: 10.1186/1742-4690-8-95. PubMed PMID: 22093708; PubMed Central PMCID:PMC3228677.

71. Valcour V, Chalermchai T, Sailasuta N, Marovich M, Lerdlum S, Suttichom D, et al. Central nervous system viral invasion and inflammation during acute HIV infection. J Infect Dis. 2012;206(2):275–82. Epub 20120502. doi: 10.1093/infdis/jis326. PubMed PMID: 22551810; PubMed Central PMCID:PMC3490695.

72. Liu Y, Tang XP, McArthur JC, Scott J, Gartner S. Analysis of human immunodeficiency virus type 1 gp160 sequences from a patient with HIV dementia: evidence for monocyte trafficking into brain. J Neurovirol. 2000;6 Suppl 1:S70–81. PubMed PMID: 10871768.

73. Suzuki K, Zaunders J, Gates TM, Levert A, Butterly S, Liu Z, et al. Elevation of cell-associated HIV-1 transcripts in CSF CD4+ T cells, despite effective antiretroviral therapy, is linked to brain injury. Proc Natl Acad Sci U S A. 2022;119(48):e2210584119. Epub 20221121. doi: 10.1073/pnas.2210584119. PubMed PMID: 36413502; PubMed Central PMCID:PMC9860316.

74. Zayyad Z, Spudich S. Neuropathogenesis of HIV: from initial neuroinvasion to HIV-associated neurocognitive disorder (HAND). Curr HIV/AIDS Rep. 2015;12(1):16–24. doi: 10.1007/s11904-014-0255-3. PubMed PMID: 25604237; PubMed Central PMCID:PMC4741099.

75. Andras IE, Pu H, Deli MA, Nath A, Hennig B, Toborek M. HIV-1 Tat protein alters tight junction protein expression and distribution in cultured brain endothelial cells. J Neurosci Res. 2003;74(2):255–65. doi: 10.1002/jnr.10762. PubMed PMID: 14515355.

76. Louboutin JP, Agrawal L, Reyes BA, Van Bockstaele EJ, Strayer DS. HIV-1 gp120-induced injury to the blood-brain barrier: role of metalloproteinases 2 and 9 and relationship to oxidative stress. J Neuropathol Exp Neurol. 2010;69(8):801–16. doi: 10.1097/NEN.0b013e3181e8c96f. PubMed PMID: 20613638; PubMed Central PMCID:PMC4707960.

77. Nakagawa S, Castro V, Toborek M. Infection of human pericytes by HIV-1 disrupts the integrity of the blood-brain barrier. J Cell Mol Med. 2012;16(12):2950–7. doi: 10.1111/j.1582-4934.2012.01622.x. PubMed PMID: 22947176; PubMed Central PMCID:PMC3524391.

78. Hernandez C, Gorska AM, Eugenin E. Mechanisms of HIV-mediated blood-brain barrier compromise and leukocyte transmigration under the current antiretroviral era. iScience. 2024;27(3):109236. Epub 20240215. doi: 10.1016/j.isci.2024.109236. PubMed PMID: 38487019; PubMed Central PMCID:PMC10937838.

79. Micci L, Alvarez X, Iriele RI, Ortiz AM, Ryan ES, McGary CS, et al. CD4 depletion in SIV-infected macaques results in macrophage and microglia infection with rapid turnover of infected cells. PLoS Pathog. 2014;10(10):e1004467. Epub 20141030. doi: 10.1371/journal.ppat.1004467. PubMed PMID: 25356757; PubMed Central PMCID:PMC4214815.

80. Xu X, Niu M, Lamberty BG, Emanuel K, Ramachandran S, Trease AJ, et al. Microglia and macrophages alterations in the CNS during acute SIV infection: A single-cell analysis in rhesus macaques. PLoS Pathog. 2024;20(9):e1012168. Epub 20240916. doi: 10.1371/journal.ppat.1012168. PubMed PMID: 39283947; PubMed Central PMCID:PMC11426456.

81. Krivine A, Force G, Servan J, Cabee A, Rozenberg F, Dighiero L, et al. Measuring HIV-1 RNA and interferon-alpha in the cerebrospinal fluid of AIDS patients: insights into the pathogenesis of AIDS Dementia Complex. J Neurovirol. 1999;5(5):500–6. doi: 10.3109/13550289909045379. PubMed PMID: 10568887.

82. Perrella O, Carreiri PB, Perrella A, Sbreglia C, Gorga F, Guarnaccia D, et al. Transforming growth factor beta-1 and interferon-alpha in the AIDS dementia complex (ADC): possible relationship with cerebral viral load? Eur Cytokine Netw. 2001;12(1):51–5. PubMed PMID: 11282546.

83. Rho MB, Wesselingh S, Glass JD, McArthur JC, Choi S, Griffin J, et al. A potential role for interferon-alpha in the pathogenesis of HIV-associated dementia. Brain Behav Immun. 1995;9(4):366–77. doi: 10.1006/brbi.1995.1034. PubMed PMID: 8903853.

84. Thaney VE, Kaul M. Type I Interferons in NeuroHIV. Viral Immunol. 2019;32(1):7–14. Epub 20180927. doi: 10.1089/vim.2018.0085. PubMed PMID: 30260742; PubMed Central PMCID:PMC6350057.

85. Anderson AM, Lennox JL, Mulligan MM, Loring DW, Zetterberg H, Blennow K, et al. Cerebrospinal fluid interferon alpha levels correlate with neurocognitive impairment in ambulatory HIV-Infected individuals. J Neurovirol. 2017;23(1):106–12. Epub 20160711. doi: 10.1007/s13365-016-0466-z. PubMed PMID: 27400930; PubMed Central PMCID:PMC5226942.

86. Boreland AJ, Stillitano AC, Lin HC, Abbo Y, Hart RP, Jiang P, et al. Sustained type I interferon signaling after human immunodeficiency virus type 1 infection of human iPSC derived microglia and cerebral organoids. iScience. 2024;27(5):109628. Epub 20240328. doi: 10.1016/j.isci.2024.109628. PubMed PMID: 38628961; PubMed Central PMCID:PMC11019286.

87. Akiyama H, Jalloh S, Park S, Lei M, Mostoslavsky G, Gummuluru S. Expression of HIV-1 Intron-Containing RNA in Microglia Induces Inflammatory Responses. J Virol. 2021;95(5). Epub 20201209. doi: 10.1128/JVI.01386-20. PubMed PMID: 33298546; PubMed Central PMCID:PMC8092841.

88. Min AK, Javidfar B, Missall R, Doanman D, Durens M, Graziani M, et al. HIV-1 infection of genetically engineered iPSC-derived central nervous system-engrafted microglia in a humanized mouse model. J Virol. 2023;97(12):e0159523. Epub 20231130. doi: 10.1128/jvi.01595-23. PubMed PMID: 38032195; PubMed Central PMCID:PMC10734545.

89. Ryan SK, Gonzalez MV, Garifallou JP, Bennett FC, Williams KS, Sotuyo NP, et al. Neuroinflammation and EIF2 Signaling Persist despite Antiretroviral Treatment in an hiPSC Tri-culture Model of HIV Infection. Stem Cell Reports. 2020;14(5):991. doi: 10.1016/j.stemcr.2020.04.006. PubMed PMID: 32402270; PubMed Central PMCID:PMC7221088.

90. Krasemann S, Madore C, Cialic R, Baufeld C, Calcagno N, El Fatimy R, et al. The TREM2-APOE Pathway Drives the Transcriptional Phenotype of Dysfunctional Microglia in Neurodegenerative Diseases. Immunity. 2017;47(3):566–81 e9. doi: 10.1016/j.immuni.2017.08.008. PubMed PMID: 28930663; PubMed Central PMCID:PMC5719893.

91. Perry VH, Crocker PR, Gordon S. The blood-brain barrier regulates the expression of a macrophage sialic acid-binding receptor on microglia. J Cell Sci. 1992;101 ( Pt 1):201–7. doi: 10.1242/jcs.101.1.201. PubMed PMID: 1569124.

92. Ostendorf L, Dittert P, Biesen R, Duchow A, Stiglbauer V, Ruprecht K, et al. SIGLEC1 (CD169): a marker of active neuroinflammation in the brain but not in the blood of multiple sclerosis patients. Sci Rep. 2021;11(1):10299. Epub 20210513. doi: 10.1038/s41598-021-89786-0. PubMed PMID: 33986412; PubMed Central PMCID:PMC8119413.

93. Groh J, Ribechini E, Stadler D, Schilling T, Lutz MB, Martini R. Sialoadhesin promotes neuroinflammation-related disease progression in two mouse models of CLN disease. Glia. 2016;64(5):792–809. Epub 20160117. doi: 10.1002/glia.22962. PubMed PMID: 26775238.

94. Gutierrez-Martinez E, Benet Garrabe S, Mateos N, Erkizia I, Nieto-Garai JA, Lorizate M, et al. Actin-regulated Siglec-1 nanoclustering influences HIV-1 capture and virus-containing compartment formation in dendritic cells. Elife. 2023;12. Epub 20230320. doi: 10.7554/eLife.78836. PubMed PMID: 36940134; PubMed Central PMCID:PMC10065798.

95. Galao RP, Pickering S, Curnock R, Neil SJ. Retroviral retention activates a Syk-dependent HemITAM in human tetherin. Cell Host Microbe. 2014;16(3):291–303. doi: 10.1016/j.chom.2014.08.005. PubMed PMID: 25211072; PubMed Central PMCID:PMC4161388.

96. Krapp C, Hotter D, Gawanbacht A, McLaren PJ, Kluge SF, Sturzel CM, et al. Guanylate Binding Protein (GBP) 5 Is an Interferon-Inducible Inhibitor of HIV-1 Infectivity. Cell Host Microbe. 2016;19(4):504–14. Epub 20160317. doi: 10.1016/j.chom.2016.02.019. PubMed PMID: 26996307.

97. Dick AD, Pell M, Brew BJ, Foulcher E, Sedgwick JD. Direct ex vivo flow cytometric analysis of human microglial cell CD4 expression: examination of central nervous system biopsy specimens from HIV-seropositive patients and patients with other neurological disease. AIDS. 1997;11(14):1699–708. doi: 10.1097/00002030-199714000-00006. PubMed PMID: 9386804.

98. Wang J, Crawford K, Yuan M, Wang H, Gorry PR, Gabuzda D. Regulation of CC chemokine receptor 5 and CD4 expression and human immunodeficiency virus type 1 replication in human macrophages and microglia by T helper type 2 cytokines. J Infect Dis. 2002;185(7):885–97. Epub 20020311. doi: 10.1086/339522. PubMed PMID: 11920312.

99. Pertel T, Hausmann S, Morger D, Zuger S, Guerra J, Lascano J, et al. TRIM5 is an innate immune sensor for the retrovirus capsid lattice. Nature. 2011;472(7343):361–5. doi: 10.1038/nature09976. PubMed PMID: 21512573; PubMed Central PMCID:PMC3081621.

100. Postler TS, Desrosiers RC. The cytoplasmic domain of the HIV-1 glycoprotein gp41 induces NF-kappaB activation through TGF-beta-activated kinase 1. Cell Host Microbe. 2012;11(2):181–93. doi: 10.1016/j.chom.2011.12.005. PubMed PMID: 22341466; PubMed Central PMCID:PMC3285415.

101. Berg RK, Melchjorsen J, Rintahaka J, Diget E, Soby S, Horan KA, et al. Genomic HIV RNA induces innate immune responses through RIG-I-dependent sensing of secondary-structured RNA. PLoS One. 2012;7(1):e29291. Epub 20120103. doi: 10.1371/journal.pone.0029291. PubMed PMID: 22235281; PubMed Central PMCID:PMC3250430.

102. Solis M, Nakhaei P, Jalalirad M, Lacoste J, Douville R, Arguello M, et al. RIG-I-mediated antiviral signaling is inhibited in HIV-1 infection by a protease-mediated sequestration of RIG-I. J Virol. 2011;85(3):1224–36. Epub 20101117. doi: 10.1128/JVI.01635-10. PubMed PMID: 21084468; PubMed Central PMCID:PMC3020501.

103. Gao D, Wu J, Wu YT, Du F, Aroh C, Yan N, et al. Cyclic GMP-AMP synthase is an innate immune sensor of HIV and other retroviruses. Science. 2013;341(6148):903–6. Epub 20130808. doi: 10.1126/science.1240933. PubMed PMID: 23929945; PubMed Central PMCID:PMC3860819.

104. Lahaye X, Satoh T, Gentili M, Cerboni S, Conrad C, Hurbain I, et al. The capsids of HIV-1 and HIV-2 determine immune detection of the viral cDNA by the innate sensor cGAS in dendritic cells. Immunity. 2013;39(6):1132–42. Epub 20131121. doi: 10.1016/j.immuni.2013.11.002. PubMed PMID: 24269171.

105. Vermeire J, Roesch F, Sauter D, Rua R, Hotter D, Van Nuffel A, et al. HIV Triggers a cGAS-Dependent, Vpu- and Vpr-Regulated Type I Interferon Response in CD4(+) T Cells. Cell Rep. 2016;17(2):413-24. doi: 10.1016/j.celrep.2016.09.023. PubMed PMID: 27705790.

106. Beignon AS, McKenna K, Skoberne M, Manches O, DaSilva I, Kavanagh DG, et al. Endocytosis of HIV-1 activates plasmacytoid dendritic cells via Toll-like receptor-viral RNA interactions. J Clin Invest. 2005;115(11):3265–75. Epub 20051013. doi: 10.1172/JCI26032. PubMed PMID: 16224540; PubMed Central PMCID:PMC1253628.

107. Fonteneau JF, Larsson M, Beignon AS, McKenna K, Dasilva I, Amara A, et al. Human immunodeficiency virus type 1 activates plasmacytoid dendritic cells and concomitantly induces the bystander maturation of myeloid dendritic cells. J Virol. 2004;78(10):5223–32. doi: 10.1128/jvi.78.10.5223-5232.2004. PubMed PMID: 15113904; PubMed Central PMCID:PMC400371.

108. McCauley SM, Kim K, Nowosielska A, Dauphin A, Yurkovetskiy L, Diehl WE, et al. Intron-containing RNA from the HIV-1 provirus activates type I interferon and inflammatory cytokines. Nat Commun. 2018;9(1):5305. Epub 20181213. doi: 10.1038/s41467-018-07753-2. PubMed PMID: 30546110; PubMed Central PMCID:PMC6294009.

109. Guney MH, Nagalekshmi K, McCauley SM, Carbone C, Aydemir O, Luban J. IFIH1 (MDA5) is required for innate immune detection of intron-containing RNA expressed from the HIV-1 provirus. Proc Natl Acad Sci U S A. 2024;121(29):e2404349121. Epub 20240710. doi: 10.1073/pnas.2404349121. PubMed PMID: 38985764; PubMed Central PMCID:PMC11260138.

110. Cenker JJ, Stultz RD, McDonald D. Brain Microglial Cells Are Highly Susceptible to HIV-1 Infection and Spread. AIDS Res Hum Retroviruses. 2017;33(11):1155–65. Epub 20170626. doi: 10.1089/AID.2017.0004. PubMed PMID: 28486838; PubMed Central PMCID:PMC5665495.

111. Haenseler W, Sansom SN, Buchrieser J, Newey SE, Moore CS, Nicholls FJ, et al. A Highly Efficient Human Pluripotent Stem Cell Microglia Model Displays a Neuronal-Co-culture-Specific Expression Profile and Inflammatory Response. Stem Cell Reports. 2017;8(6):1727–42. doi: 10.1016/j.stemcr.2017.05.017. PubMed PMID: 28591653; PubMed Central PMCID:PMC5470330.

112. Martins-Ferreira R, Leal B, Costa PP, Ballestar E. Microglial innate memory and epigenetic reprogramming in neurological disorders. Prog Neurobiol. 2021;200:101971. Epub 20201209. doi: 10.1016/j.pneurobio.2020.101971. PubMed PMID: 33309803.

113. Munera JO, Sundaram N, Rankin SA, Hill D, Watson C, Mahe M, et al. Differentiation of Human Pluripotent Stem Cells into Colonic Organoids via Transient Activation of BMP Signaling. Cell Stem Cell. 2019;24(5):829. doi: 10.1016/j.stem.2019.04.002. PubMed PMID: 31051135; PubMed Central PMCID:PMC6530774.

114. Snoeck HW. Modeling human lung development and disease using pluripotent stem cells. Development. 2015;142(1):13–6. doi: 10.1242/dev.115469. PubMed PMID: 25516965.

